# Temporal sensitivity for achromatic and chromatic flicker across the visual cortex

**DOI:** 10.1101/2023.07.24.550403

**Authors:** Carlyn Patterson Gentile, Manuel Spitschan, Huseyin O. Taskin, Andrew S. Bock, Geoffrey K. Aguirre

## Abstract

The retinal ganglion cells (RGCs) receive different combinations of L, M, and S cone inputs and give rise to one achromatic and two chromatic post-receptoral channels. Beyond the retina, RGC outputs are subject to filtering and normalization along the geniculo-striate pathway, ultimately producing the properties of human vision. The goal of the current study was to determine temporal sensitivity across the three post-receptoral channels in subcortical and cortical regions involved in vision. We measured functional magnetic resonance imaging (MRI) responses at 7 Tesla from three participants (two males, one female) viewing a high-contrast, flickering, spatially-uniform wide field (~140°). Stimulus flicker frequency varied logarithmically between 2 and 64 Hz and targeted the L+M+S, L–M, and S–[L+M] cone combinations. These measurements were used to create temporal sensitivity functions of primary visual cortex (V1) across eccentricity, and spatially averaged responses from lateral geniculate nucleus (LGN), V2/V3, hV4, and V3A/B. Functional MRI responses reflected known properties of the visual system, including higher peak temporal sensitivity to achromatic vs. chromatic stimuli, and low-pass filtering between the LGN and V1. Peak temporal sensitivity increased across levels of the cortical visual hierarchy. Unexpectedly, peak temporal sensitivity varied little across eccentricity within area V1. Measures of adaptation and distributed pattern activity revealed a subtle influence of 64 Hz achromatic flicker in area V1, despite this stimulus evoking only a minimal overall response. Comparison of measured cortical responses to a model of integrated retinal output to our stimuli demonstrates that extensive filtering and amplification is applied to post-retinal signals.

**Significance Statement:** We report the temporal sensitivity of human visual cortex across the three canonical post-receptoral channels from central vision to the far periphery. Functional MRI measurements of responses from the LGN, V1, and higher visual cortical areas demonstrate modification of temporal sensitivity across the visual hierarchy. This includes amplification of chromatic signals between the LGN and V1, and an increase in peak temporal sensitivity in visual areas beyond V1. Within V1, we find a surprising stability of peak temporal sensitivity in the periphery for all three post-receptoral directions. Comparison of our results to a model of retinal output demonstrates the presence of substantial post-retinal filtering, yielding greater uniformity of responses across area V1 than would be predicted from unmodified retinal signals.

## Introduction

Cone and retinal ganglion cell (RGC) density declines exponentially from the fovea to the periphery. There is a corresponding and well-studied change in the spatial frequency sensitivity (acuity) of human perception that reflects this gradient (Kelly, 1984; e.g., spatial acuity; Thibos et al., 1996). Although less numerous, cell populations in the peripheral retina are found to have faster kinetics (Seiple and Holopigian, 1996; Solomon, 2005), raising the possibility that visual perception has a corresponding change in temporal frequency sensitivity with eccentricity.

Signals from the cones are combined into three post-receptoral channels (Figure 1a): one achromatic or luminance (L+M) channel, and two chromatic channels, red-green (L-M), and blue-yellow (S–[L+M]). The eccentricity dependence of temporal sensitivity has been most studied for achromatic stimuli, and generally shows an improvement in threshold contrast detection for higher frequency flicker in peripheral as compared to central vision (Kelly, 1984; Snowden and Hess, 1992). In foveal vision, chromatic channels favor slower frequencies as compared to the achromatic channel (Swanson et al., 1987). Relatively little is known regarding how eccentricity variation influences temporal sensitivity across the chromatic post-receptoral channels, although one set of measurements made in the LM plane out to 14° found little variation in temporal response (Gelfand and Horwitz, 2018).

**Figure 1.**
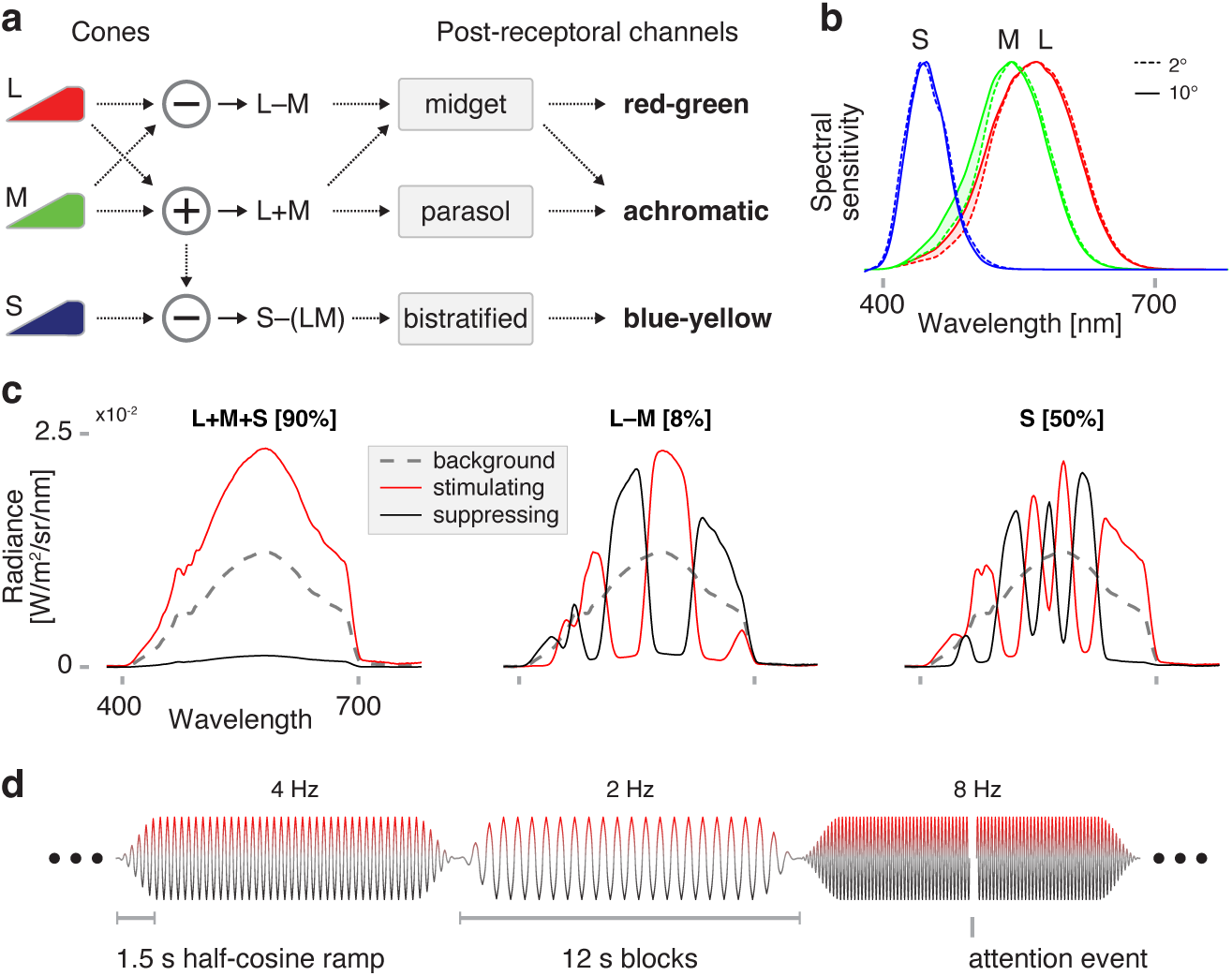
Targeting the post-receptoral channels with spectral flicker. (a) L-, M-, and S-cones interact at the level of the retina to form the achromatic (L+M), chromatic red-green (L–M), and chromatic blue-yellow (S–[L+M]) post-receptoral channels. Midget RGCs contribute substantially to both the achromatic and red-green channels. (b) The spectral sensitivities of the cones vary slightly between central (2 degrees, dotted line) and more peripheral (10 degrees, solid line) eccentricity locations, largely due to the filtering effects of macular pigment. (c) Spectral modulations were generated to target combinations of cone classes while silencing others, corresponding to the three cardinal axes of DKL space. Stimuli modulated between a suppressing (black) and stimulating (red) spectrum, around a common background (dotted gray). Different modulations produced (*left*) equal nominal stimulation of the L, M, and S cones, thus producing isolated stimulation of the L+M channel, (*center*) differential stimulation of the L and M cones, while silencing S, and (*right*) isolated stimulation of the S cones, producing targeted stimulation of the S-[L+M] channel. Nominal Michelson contrast upon the targeted post-receptoral channels is given in brackets above each plot. Spectral targeting accounted for the 2° and 10° variants of spectral sensitivity for the cone classes, to equate nominal stimulus contrast across the visual field. The stimuli presented to P3 differed in spectral form, but had the same contrast, luminance, and background chromaticity. (d) Temporal flicker was presented at logarithmically spaced frequencies between 2 Hz and 64 Hz (as well as 0 Hz) in each of many 12 second trials. The modulation was subjected to a half-cosine ramp at onset and offset. The ordering of the stimulus frequencies was specified by a pseudo-random, counterbalanced sequence. During intermittent attention events the entire stimulus field dimmed for 250 msecs, in response to which the participant was asked to make a bilateral button press. Different fMRI acquisitions presented modulations that targeted a different post-receptoral channel. Extended Figure 1-1 and Extended Table 1-1 present estimates of the intended and inadvertent contrast produced by the stimuli upon the cone photoreceptors.

The properties of threshold detection as measured with psychophysics may differ in meaningful ways from the perceptual experience of supra-threshold stimuli (i.e., natural vision). Neural responses to supra-threshold stimuli might be expected to reflect the properties of visual perception. Electrophysiological studies in non-human primates demonstrate filtering of temporal information at successive stages of the visual system; the largest decline in temporal sensitivity occurs between the cones and RGCs, with additional filtering of high-temporal frequency information between the lateral geniculate nucleus (LGN) and cortex (Hawken et al., 1996; Horwitz, 2020, 2021a). Consequently, we may expect that the temporal sensitivity of human visual cortex—and presumably human supra-threshold perception—reflects a substantial modification of the initial retinal signal.

Here we report measurements of the regional and eccentricity dependence of the temporal sensitivity of human visual cortex. We studied three participants using a 7 Tesla scanner while they viewed high-contrast, wide-field stimuli that targeted each of the three post-receptoral channels, modulating across a range of flicker frequencies. Our results provide temporal sensitivity functions across multiple brain regions representing stages of the visual pathway from lateral geniculate nucleus (LGN) to extra-striate cortex. More detailed measurements across eccentricity were also obtained for the primary visual cortex (area V1). Our findings confirm several temporal filtering properties of the visual system that have been previously reported, as well as normalization of signals in area V1 across post-receptoral channel and eccentricity relative to the retina and LGN. Unexpectedly, we find that peak temporal sensitivity is stable, and perhaps slightly declines, across eccentricity for all three post-receptoral directions.

## Materials and Methods

### Participants

The three participants are authors of this report (P1 age 46; P2 age 32; P3 age 39), the first two of whom are male. The participants had normal or corrected to normal acuity and normal color vision. This study was approved by the University of Pennsylvania Institutional Review Board in accordance with the Declaration of Helsinki, and all participants provided written consent.

### Custom contact lens

The participants were fitted with custom contact lenses that provided either +36 or +32 diopters of magnification, with the latter selected to complement the –4 diopter myopia for that participant (P1). The contact lenses were Alden Optical, HP54 spherical soft lenses, constructed from 54% Hioxifilcon D (sourced from Benz Research and Development). The lens was 0.59 mm at its thickest point. During scanning, the participant wore a contact lens in the right eye. Prior to each scanning session, the right eye underwent pharmacologic dilation to provide increased and matched retinal illuminance across the participants; the left eye was patched.

### Spectral modulations

Stimuli were generated using a multi-primary light engine. For participants P1 and P2, the stimuli were created using the OneLight Spectra device, which produces desired spectra through the use of a digital micro-mirror device chip, and was configured to mix 56 independent primaries with a FWHM of ~16 nm at a refresh rate of 256 Hz. For participant P3, the stimuli were created using a CombiLED (Prizmatix Inc) which combines the output of 8, narrow-band, fiber-coupled LEDs at a refresh rate of 400 Hz. Spectral modulations were designed to target specific photoreceptor classes alone or in combination, using the silent substitution approach (Estévez and Spekreijse, 1982). The design of these stimuli has been described in detail previously (Spitschan et al., 2015; Spitschan and Woelders, 2018; Barnett et al., 2021). Briefly, a set of three modulations were created, which targeted L–M, S, and LMS cone combinations; the last of these was a “light flux” modulation, in which the entire stimulus spectrum was scaled. All three stimulus types were modulations around a common background.

The chromatic L–M and S modulations were designed to minimize differences in contrast between the fovea and the periphery. This was achieved by treating the cone fundamentals at 2 and 10 degrees (CIE, 2006) as different classes of photoreceptors, and tailoring modulations to produce equivalent contrast on these two classes (Barnett et al., 2021). The cone fundamentals were adjusted for age-related differences in lens density (CIE, 2006), and the transmittance spectrum of the contact lens material was measured and incorporated into the model of pre-receptoral filtering.

Spectroradiometric measurements of the stimuli were made before each experimental session (PhotoResearch PR-670). The nominal maximum contrast of the modulation was approximately 90% for LMS, 50% for S, and 8% for L–M. Extended Figure 1-1 and Extended Table 1-1 provide the predicted contrast of the stimuli as a function of eccentricity and in the presence of imprecision in device control. The chromaticity of the background was x = 0.4025, y = 0.4486, luminance was Y = ~400 cd/m^2^; chromaticity and luminance computed with respect to the XYZ 10° physiologically relevant color matching functions.

The stimuli were conveyed via a fiber-optic cable to a custom-made, MR-compatible eye piece that was positioned over the right eye of the participant. After passing through ultra-violet and heat-absorbing glass, light was back-projected on a 35 mm diameter, opal diffusing glass. The stimulus surface was positioned ~10 mm from the eye. This geometry provided a uniform stimulus field of ~60° radial eccentricity. The +36 diopter contact lens produced approximately x1.18 spectacle magnification. The resulting stimulus field was therefore close to 70° radial eccentricity.

### Scanning environment and MR imaging of the brain

An MR imaging session was conducted for each participant on each of two days within the same week. During the scanning session, all sources of light within the scanner room were extinguished and covered. Room lights were on when participants were positioned within the scanner, but these were turned off at the start of data acquisition.

All MRI images were obtained with a 32-channel head coil on a 7T Siemens scanner. The anatomical T1w images were acquired with voxel size=0.8 mm iso, resolution=480×640×176, TR = 2800ms, TE=4.28ms, TI=1500ms, and flip angle=7°. For the BOLD images, a multiband epfid2d1_130 sequence was used with multiband acceleration factor=5, voxel size=1.6 mm iso, resolution=130×130×75, TR=1000ms, TE=26.8ms, flip angle=45°, and average=1.

### Experimental design

Each fMRI acquisition was 336 seconds in duration and presented modulations that targeted one particular post-receptoral channel (LMS, L–M, or S). The acquisition was composed of 28, 12 second trials. Each trial presented a spectral modulation that flickered at one of 7, log-spaced frequencies (0, 2, 4, 8, 16, 32, 64 Hz). A pseudo-random, counterbalanced (Aguirre et al., 2011) ordering of the frequencies was defined (50 elements). This sequence was divided into two-subsequences (“A” and “B”), each padded with three instances of the 0 Hz trial at the end. A set of 6 acquisitions composed a scanning “block”, and consisted of the ordered acquisitions: LMS_A_, L–M_A_, S_A_, S_B_, L–M_B_, LMS_B_. A total of 6 blocks were collected for each participant across the two sessions. The stimulus field was set to the shared spectral background before and between each acquisition.

The sinusoidal spectral modulation presented in each block was windowed by a 1.5 second, half-cosine ramp at trial onset and offset. An attention task was designed to confirm that participants were awake and monitoring the stimuli, but minimally interfere with the measurement of the response to the flicker stimuli. The attention event was a 250 ms darkening of the stimulus field that occurred with 33% probability during each trial. The event could appear at any point during the trial, except constrained not to take place during first or last 2 seconds of the trial. The participant was asked to make a bilateral button press on a fiber-optic pad in response to each attention event. Correct detection of the >300 attention events was high for the three participants (P1 98%; P2 94%; P3 98%), and false alarm rates were all below 0.5%.

### Pre-processing of fMRI data

Three-dimensional surface reconstruction of the anatomical images for both participants was performed using Freesurfer (Dale et al., 1999). For the preprocessing of the BOLD images, we used the *fmriprep* pipeline with custom options (Esteban et al., 2019). The pipeline included bias field correction (Tustison et al., 2010), brain extraction and normalization (Avants et al., 2008), tissue segmentation (Zhang et al., 2001), slice timing correction (Cox, 1996), and motion correction (Jenkinson et al., 2002). As no B0 field maps were collected, we performed susceptibility correction with symmetric group-wise diffeomorphic normalization (*SyN*). This method aims to correct distortion by inverting the intensity of BOLD images (Huntenburg, 2014; Wang et al., 2017) and registering these inverted images to subject T1w image using an average fieldmap template as the constraint (Treiber et al., 2016). Noise regressors were obtained with AROMA (Pruim et al., 2015). The set of regressors was inspected and any component with a spatial extent that was largely confined to the occipital cortex was excluded. The effect of the remaining set of regressors was removed from the data using the “non-aggressive” denoising strategy. Finally, we mapped the preprocessed volumetric BOLD data to HCP CIFTI FSLR_32k template space with *ciftify* which uses the algorithms and templates from the HCP minimal processing pipeline (Glasser et al., 2013) to map volumetric timeseries data to CIFTI surfaces.

### Modeling of fMRI time-series data

The time-series data in each voxel was fit with a non-linear model that simultaneously estimated the amplitudes of response to the stimuli and the shape of the hemodynamic response function (similar to Prince et al., 2022). First, the time-series data in each voxel in each acquisition were converted to percentage change units. The data from all acquisitions across both sessions were then concatenated, for a total of 12,096 time points. The amplitude of response to each stimulus frequency was modeled as a 12 second step function. Attention events were modeled as delta functions. All trials of a given stimulus frequency in a single acquisition were modeled with a single covariate, but different covariates modeled the response in each acquisition (for a total of 8×36 = 288 covariates). The covariates were subject to convolution by a model of the hemodynamic response function, which itself was under the control of three parameters that specify the weighted combination of an orthogonal basis (Woolrich et al., 2004). The form of the HRF was held in common across all acquisitions for a given voxel for a given participant. Within a non-linear model fitting routine the shape of the HRF and the amplitudes of the covariates were adjusted to minimize the L2 norm of the difference between the observed and modeled fMRI time series, after subjecting the model and data to a 0.0387 Hz high-pass filter. Because this analysis was computationally demanding and thus time consuming, we confined the analysis to the posterior 1/3^rd^ of the cortex, which included all retinotopically organized early visual areas.

This analysis yielded estimates of the amplitude of neural response to each stimulus frequency for each acquisition, providing 12 measures in total for each crossing of post-receptoral channel x stimulus frequency for each participant. For display of the fit of the model to the time-series data, and for voxel selection, we first regressed out the effect of the attention task, and then averaged across the six acquisitions for a given post-receptoral direction with stimulus ordering “A”, and the six acquisitions with stimulus ordering “B”, and concatenated these. The R^2^ value of the model fit to these average data (which excludes variance attributable to the attention task) was retained.

### Anatomical regions of interest

An anatomic model of the retinotopic organization of visual cortex was used to define the location and eccentricity mapping of several early visual areas for each participant (Benson et al., 2014, 2018). For the analysis across eccentricity, the V1 ROI was divided into six, log-spaced divisions of eccentricity (division edges: 0, 2.8, 5.6, 11.25, 22.5, 45, and 90°). The LGN ROI was defined in CIFTI FSLR_32k space. We segmented the LGN using the Freesurfer thalamic segmentation algorithm (Iglesias et al., 2018). A warp was calculated between participant anatomy and the Montreal Neurological Institute (MNI) space, and the segmentation masks were moved to the MNI coordinates with nearest neighbor interpolation. The reason for this operation is because the CIFTI FSLR atlas uses MNI for the subcortical voxel maps. Within a given eccentricity band (or region mask), voxels with a time-series model fit of R^2^ > 0.1 were selected and averaged to obtain temporal sensitivity measurements.

### Modeling of the temporal sensitivity function

TSFs were fit with a difference-of-exponentials model (Hawken et al., 1996) developed from Watson’s temporal sensitivity model (Watson, 1986):

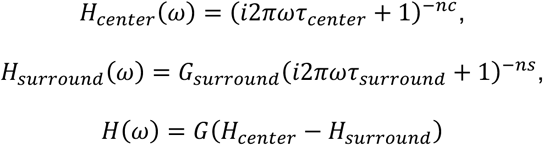

where *nc* and *ns* represent the order of the center (fast) and surround (slow) filters, fixed at 9 and 10, respectively (Watson, 1986). Variables that were free to vary included *τ_center_*, the multiplier of the *τ_surround_* relative to the *τ_center_*, and the multiplier of the *H_surround_* relative to the *H_center_*. Linear constraints were placed on the model parameters, and an additional non-linear constraint was added to ensure the model produced a unimodal fit. The amplitudes of BOLD fMRI response across temporal frequency were fit with this model using the Matlab *fmincon* routine. While the Watson model provided a good description of the temporal sensitivity functions in our data, we found the model to be over-parameterized, in that model fits did not provide unique parameter solutions. Consequently, the values of the parameters were not especially informative. We instead summarized the model fits by noting the amplitude and temporal frequency at which the maximum interpolated response was found.

### Statistical analysis

Goodness-of-fit for fMRI timeseries modeling was determined by R^2^. TSFs are shown with the median and interquartile range (IQR). Median and IQR amplitude and peak temporal frequency were determined by fitting 1,000 TSFs bootstrapped across acquisitions under each stimulus condition. A map of temporal sensitivity was also created. To do so, we fit the Watson temporal sensitivity function to the response data from each vertex, separately for each stimulus direction. At each vertex, we found the interpolated temporal frequency corresponding to the maximum response (expressed as log Hz). The set of frequency values across vertices was then z-transformed by subtracting the across-voxel mean and then dividing by the standard deviation. The three z-maps obtained from the three stimulus directions were then averaged to provide an overall map of relative peak temporal sensitivity.

An analysis of the carry-over effects of stimulus frequency was performed (Aguirre, 2007). First, the residual fMRI signal from area V1 was obtained after fitting the time-series data to recover the amplitude of response to each stimulus type. Next, we constructed a regression matrix that modeled the effect of presentation of each stimulus frequency upon the response to every other stimulus frequency, including itself. Given the seven different frequencies studied (including the 0Hz condition), this resulted in a set of 49 covariates. These covariates were generated separately for trials that presented the LMS modulation, and for trials that presented a chromatic modulation (either L–M or S directed). After convolution of the covariates with the subject-specific HRF measured for area V1, we regressed the residual time-series data upon the set of covariates and obtained the modulatory effect of each stimulus upon every other stimulus. These values were arranged into a carry-over matrix for presentation.

We conducted an analysis of the multi-vertex pattern of responses within area. We obtained the amplitude of response during each fMRI acquisition for each stimulus (relative to the 0Hz condition) at each vertex within area V1. Within each acquisition, the set of amplitude values across vertices for each stimulus frequency was standardized to have a mean of zero and unit standard deviation. Using a split-half approach, the pattern of responses across vertices was obtained for a given stimulus frequency, and the Pearson correlation of this pattern measured against the patterns measured in the held-out data for each of the seven stimulus frequencies. This produced a pattern similarity matrix, and the average of many such matrices was obtained across all possible split-half partitions of the acquisitions.

### Model of retinal signals

We created a model of predicted retinal output to our stimuli, based upon previous *in vivo* measurements in the macaque from three classes of retinal ganglion cells (parasol, midget, small bistratified). These data were extracted from Figures 1a, b of Solomon and colleagues (2002) and Figures 9a, b and 13a of Solomon (2005). The electrophysiologic data were recorded in response to stimuli similar to those employed in our study (spatially uniform, 4.7° diameter modulations that targeted the DKL cardinal axes around a 2000 Troland background). Measurements were made for log-spaced temporal frequencies and contrasts from cells in three different retinal eccentricity bands (0-10°, 20-30°, >30°; only the first two eccentricity bands available for the bistratified responses). We averaged the “on” and “off” cell responses where both were provided. Amplitude was given in units of “gain”, defined as spikes per second per percent contrast of the stimulus relative to the maximum possible cone contrast available for a given stimulus modulation in cone contrast space (measured at the initial slope of the contrast response function).

To obtain the predicted response to our stimuli, we calculated the spikes / sec response that would be expected for the contrast level used for our stimuli, with the assumption that parasol cells saturate at 25% contrast (Lee et al., 1990). The response to the LMS modulation was taken as the sum of the midget and parasol achromatic responses. We assigned the RGC measurements to three eccentricity locations within the cell recoding bins (5°, 25°, and 40°). To produce the predicted retinal response integrated across cell populations, we scaled responses at each eccentricity by the total quantity of RGC receptive fields (Watson, 2014), the relative proportions of the different RGC classes (Dacey, 1993; Drasdo et al., 2007), and the change in annular retinal area with eccentricity. We characterized the model output by the amplitude and temporal frequency at the peak response.

### Code Accessibility

All software used for data processing and data analysis is publicly accessible. The software used for overall data analysis and figure generation is found here: https://github.com/gkaguirrelab/Patterson_2024_JNeurosci. The model of retinal response may be found here: https://github.com/gkaguirrelab/rgcTemporalSensitivity.

## Results

We obtained BOLD fMRI data from three participants (P1, P2, and P3) while they viewed a spatially uniform, wide field of flickering light. Our stimuli separately targeted the three post-receptoral channels (Figure 1a). The design of these stimuli accounted for variation in receptoral spectral sensitivity across retinal eccentricity (Figure 1b), and were designed to produce spatially uniform, high contrast on the targeted post-receptoral channel across the visual field (Figure 1c). Below we refer to these stimuli as being oriented in different “directions” along the cardinal axes of Derrington-Krauskopf-Lennie (DKL) space (Derrington et al., 1984). Across many fMRI acquisitions, these spectral modulations were flickered at log-spaced temporal frequencies ranging from 2 to 64 Hz (Figure 1d). An additional, “0 Hz” condition presented the steady stimulus background. Participants viewed the stimulus field within an MRI-compatible eye piece through a +36 or +32 D contact lens, increasing the effective stimulus radius to ~70° eccentricity. Estimates of the intended and inadvertent stimulus contrasts that reached the photoreceptors are provided in Figure 1-1 and Table 1-1.

### Variation in flicker frequency induces consistent variation in BOLD fMRI responses

First, we fit the BOLD fMRI data from each participant with a time-series model to recover the amplitude of response to each stimulus type at each voxel or vertex. The average amplitude of BOLD fMRI signal was estimated separately for each stimulus direction (LMS, L–M, S) for each stimulus frequency across the 12 second periods of stimulation; the amplitude of response to the attention events was also estimated. These estimates were made for each acquisition to allow assessment of measurement error. Figure 2a presents maps of the R^2^ fit of the time-series model across the cortical surface for the three participants, not including the effect of the attention events. The highest-quality model fits were found within the borders of area V1.

**Figure 2.**
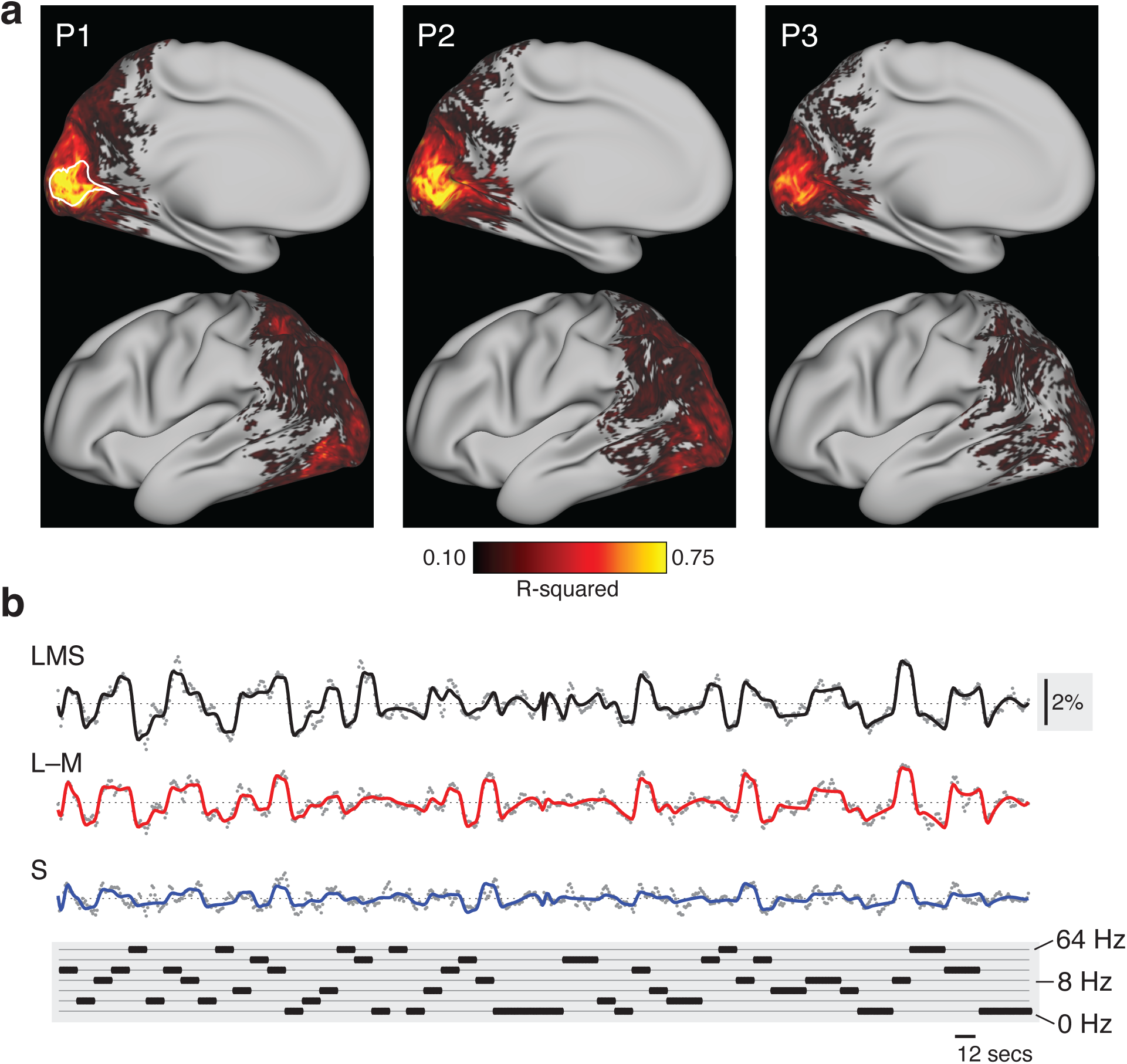
Fit to fMRI time series data. The time series data at each voxel and vertex were analyzed separately for each participant across all acquisitions. (a) The partially-inflated cortical surface map with the R^2^ value of the model fit to the time series data at each vertex, averaged across the left and right hemispheres into a single, pseudo-hemisphere. The border of area V1 is indicated with a white line on the map from participant 1. (b) The average BOLD fMRI time series for P1 (gray dots) from area V1, and across the acquisitions for each stimulus type (LMS, L–M, and S). The continuous line shows the fit of the time-series model. An inset, vertical scale bar on the right indicates a 2% BOLD fMRI signal response. An inset scale bar at bottom-right indicates a 12-second interval. The plot at the bottom provides the ordering of stimulus frequency. Inspection of the time series reveals a systematic relationship with the stimulus sequence. The time-series data for the other participants are shown in Extended Figure 2-1.

We obtained the V1 time-series data and model fit (averaged across vertices and acquisitions) for each participant (after removing the effect of the attention events) for each of the three stimulus directions. Time-series data for P1 is shown (Figure 2b), and very similar fits were found for all three participants (Figure 2-1). The analysis provided a good description of the variation in the BOLD fMRI signal. Further, the responses were systematically related to the pattern of stimulus temporal frequencies. There was clear variation in the amplitude of the cortical response as a function of the modulation frequency of the stimulus for each post-receptoral direction.

### Temporal filtering and amplification of chromatic signals between LGN and V1

The first level analysis provided the amplitude of BOLD fMRI response to each stimulus frequency. We next created temporal sensitivity functions (TSFs) which describe the average response to each stimulation frequency (relative to the 0 Hz condition). Figure 3a presents the TSFs measured for participant P1 within the LGN and area V1; P2 and P3 had similar responses (Figure 3-1). The data were fit with the Watson (1986) difference-of-exponentials temporal model, which provided a reasonable description. Visual inspection of the TSFs reveals two notable differences between LGN and V1 responses: (1) there was a relative increase in the amplitude of response to chromatic stimuli between LGN and V1; and (2) the peak temporal sensitivity shifted to lower frequencies between LGN and V1 across all post-receptoral channels. We quantified these impressions for all three participants by obtaining the median (across bootstrapped acquisitions) peak amplitude and peak temporal sensitivity provided by the fit of the Watson model (Figure 3b). Equal or greater responses were seen in V1 as compared to the LGN for all three post-receptoral channels, with a tendency for a greater increase in responses to chromatic signals between the LGN and V1. There was also a shift of peak temporal sensitivity towards lower frequencies between LGN and V1 for all three post-receptoral channels. This was largest for the achromatic stimuli, for which peak sensitivity dropped by 5-10 Hz between the thalamic and cortical sites.

**Figure 3.**
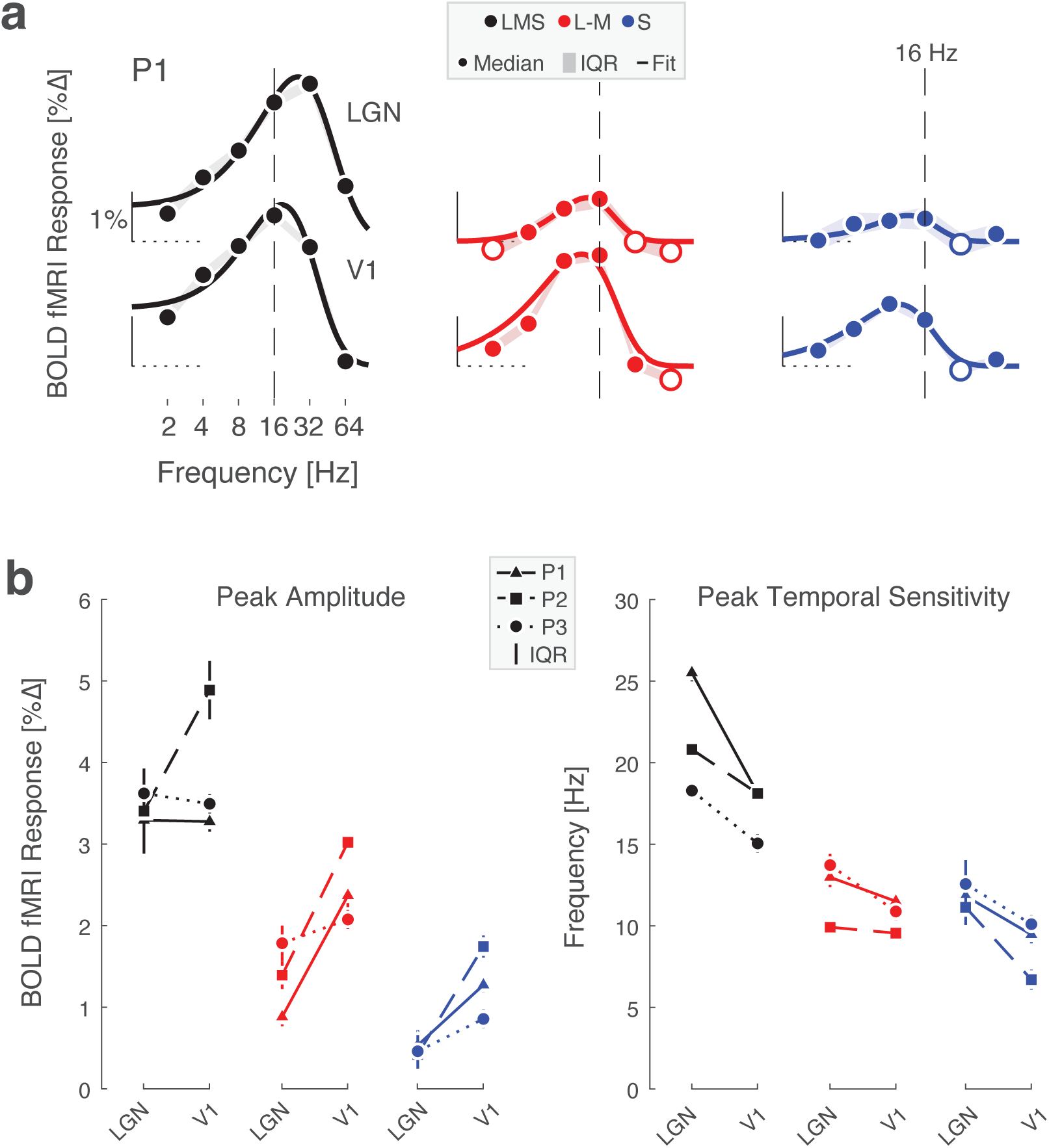
Temporal sensitivity to flicker in LGN and V1. (a) LGN and V1 TSFs for stimuli that target the 3 post-receptoral channels are shown for P1, fit with the Watson difference-of-exponentials model. Data points represent the boot-strapped median across acquisitions, and shading provides the interquartile range. Open circles indicate points where the response was less than zero. A dashed reference line is positioned at 16 Hz in each plot, facilitating a comparison of the point of peak temporal sensitivity across plots. TSFs for P2 and P3 are shown in Extended Figure 3-1. (b) Peak amplitude (left) and peak temporal sensitivity (right) for each participant across region, derived from Watson model fits. Data points represent median value and error bars represent interquartile range.

### Peak temporal sensitivity in V1 slows slightly towards the periphery

The retinotopic organization of V1 allowed us to examine temporal sensitivity across eccentricity. BOLD fMRI responses at each vertex within V1 were fit with the Watson model to obtain the peak amplitude and peak temporal frequency of the TSF. We then examined how these properties varied as a function of eccentricity (Figure 4a). The interpretation of the absolute amplitude of response is complicated by the effect of non-neural influences upon the BOLD fMRI response; we nonetheless can examine the relative response between the stimulus directions. The response to the L–M modulation is comparable to the LMS-driven response out to ~5° eccentricity, and then drops steadily into the visual periphery. In contrast, the response to S-directed stimuli across eccentricity follows the same general form as the response to LMS, although is scaled down and reaches zero close to the fovea.

**Figure 4.**
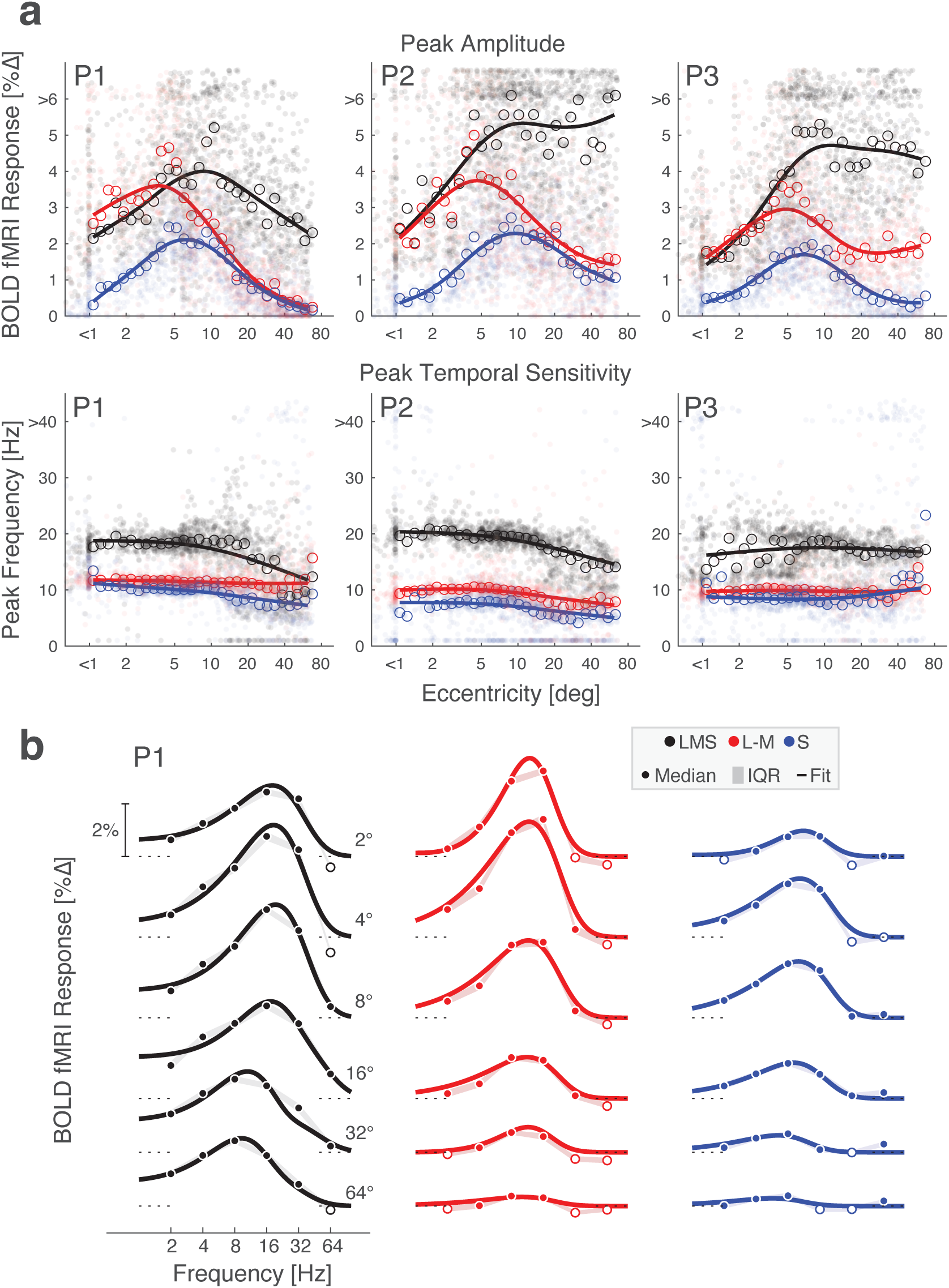
Temporal flicker sensitivity within area V1 across eccentricity. (a) Peak amplitude (top row) and peak temporal frequency (bottom row) within area V1 for P1, P2, and P3 for stimuli that targeted each of the three post-receptoral directions. Dots represent values for each vertex and lines represent the fit of a smoothing spline to the median value (open circles) at each of 30 eccentricity bins. (b) V1 TSFs at logarithmically spaced eccentricities across the three post-receptoral channels are shown for P1 with Watson temporal model fits; P2 and P3 had similar TSFs across frequency and are shown in Figure 4-1. Data points represent the boot-strapped median across acquisitions, shading provides the interquartile range, and the line provides the Watson model fit. Open circles indicate points where the response was less than zero. An additional analysis used the retinotopic assignment at each vertex in V1 to construct a map of peak temporal sensitivity in the visual field for each participant and post-receptoral direction.

We examined peak temporal sensitivity across post-receptoral channel and eccentricity. Higher peak temporal frequency values were found for the achromatic as compared to the chromatic stimuli. Peak temporal sensitivity was notably stable across eccentricity, although with some tendency for the peak response to shift towards lower frequencies starting at 10 – 20° eccentricity. With the exception of a few measurements in the very far periphery, we found no evidence of a systematic shift to higher temporal sensitivity with greater eccentricity.

Next, we divided area V1 into six concentric bands to visualize TSFs as a function of visual field eccentricity. To account for cortical magnification of the visual field, the borders of these bands were logarithmically spaced, yielding patches of cortex roughly matched in surface area. Within each band, we obtained the TSF for each stimulus modulation, which was similar for P1 (Figure 4b), P2, and P3 (Figure 4-1). Reliable TSFs were measured for all participants, for all three stimulus types, across eccentricity. This includes in the most peripheral eccentricity band (centered at 64°). Examination of these TSFs confirms the impression that the peak response to LMS stimuli tends to shift to lower temporal frequencies at greater eccentricities within area V1. This is reflected in both the peak temporal frequency, as well as in a relatively larger low-pass component to the response. To examine temporal sensitivity as a function of polar angle and eccentricity we projected the V1 responses to a visual field representation, using the retinotopic organization of area V1 inferred from cortical topology (Figure 4-2). The only systematic variation seen in this analysis was as a function of eccentricity.

### Peak temporal sensitivity increases beyond area V1

We created a map of peak temporal frequency across the cortex, combining normalized measurements from each stimulus direction (Figure 5a). This map shows the temporal sensitivity of a cortical location relative to the other posterior cortical locations. There was a relatively uniform set of values within area V1, positioned in the mid-range of relative temporal sensitivity. Within retinotopically organized extra-striate cortex, peak sensitivity shifted to higher temporal frequencies. Finally, at more anterior points within visually responsive cortex, peak sensitivity was found at lower temporal frequencies.

**Figure 5.**
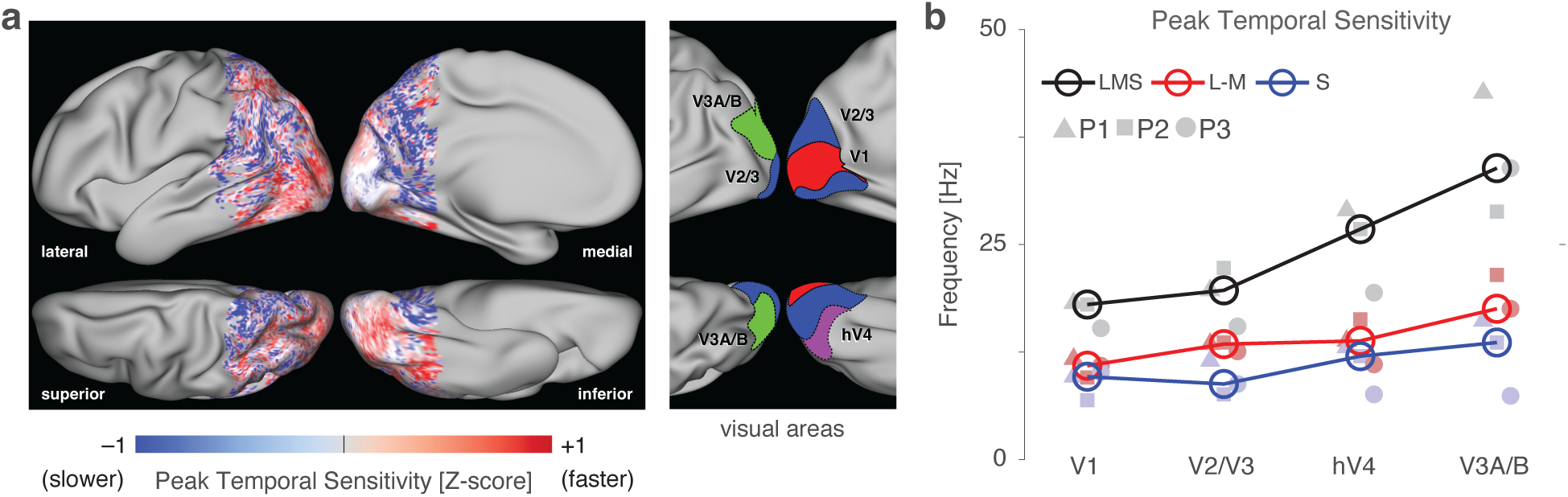
Peak temporal frequency beyond area V1. (a) Relative peak temporal sensitivity across posterior cortex, combined across stimulus directions and then averaged across the three participants. Lateral, medial, superior, and inferior views are provided. Map values reflect the relative peak frequency across the cortex, averaged across the stimulus directions. The locations of several early visual areas are indicated inset at right. (b) Median temporal sensitivity across participants (open circles) for each of several visual cortex regions of interest across the three post-receptoral channels. Values for individual participants are given by the filled plot symbols.

We measured the peak temporal sensitivity of specific extra-striate regions (Figure 5b). Peak temporal sensitivity increased across the visual hierarchy, rising slightly between V1 and V2/V3, and again into both hV4 and V3A/B. This increase was most marked in the response to the LMS stimuli.

### Effect of flicker frequency upon subsequent neural response

Presentation of many seconds of rapid, spatially uniform flicker leads to a reduction in the response of LGN neurons to subsequent stimuli that can last tens of seconds, especially for achromatic signals carried by the parasol cells (Solomon et al., 2004). We tested for these effects in our data by examining how the response to each 12s trial varied as a function of the immediately preceding stimulus. Figure 6 shows the resulting carry-over matrix (Aguirre, 2007), averaged across participants separately for the achromatic and chromatic stimuli. Each cell indicates the degree to which the fMRI response to a particular stimulus was larger (red) or smaller (black) depending upon the flicker frequency of the prior stimulus.

**Figure 6.**
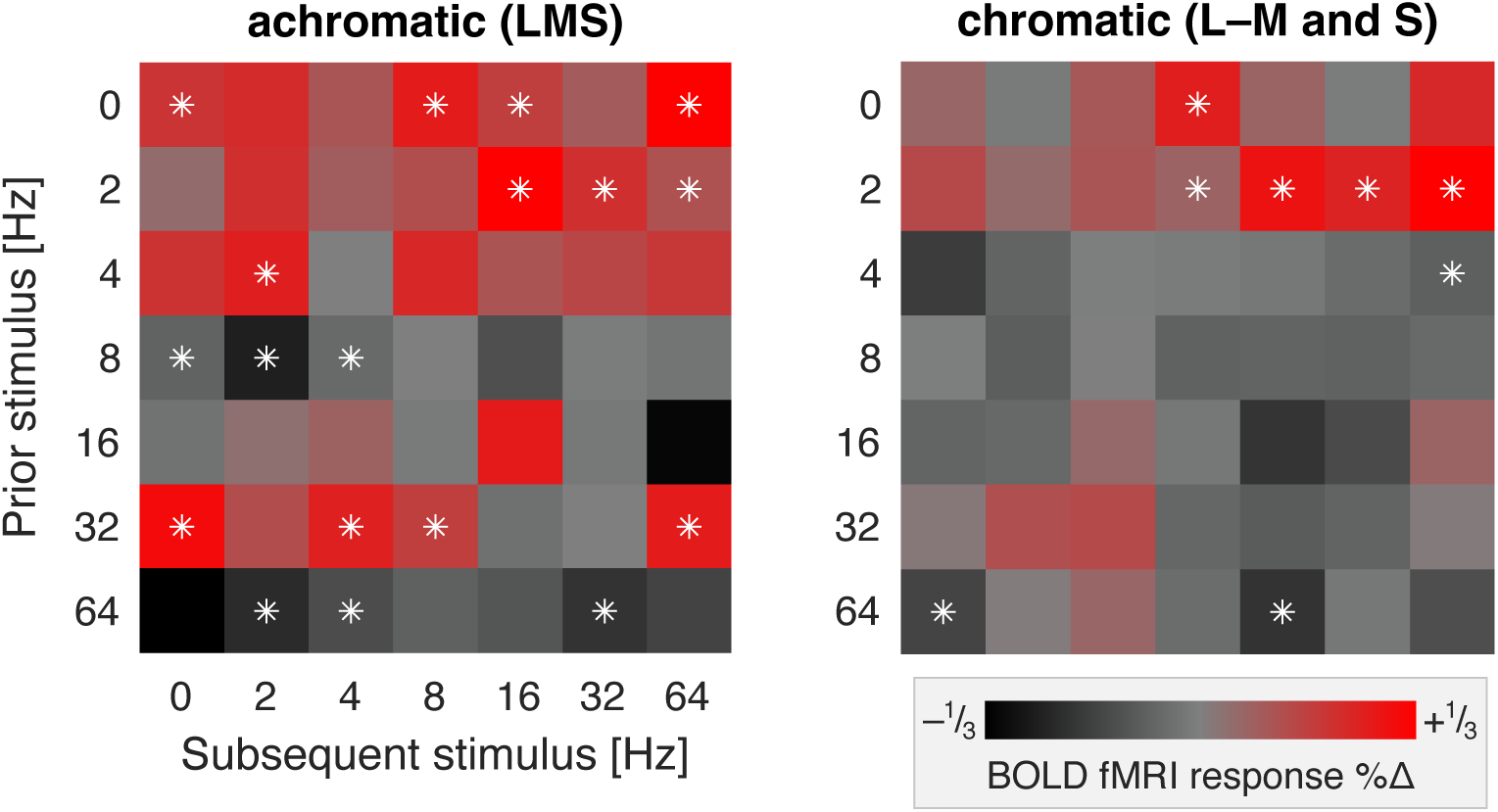
Carry-over adaptation effects in V1. Each matrix presents the modulatory effect of a given frequency of flicker stimulation (the “prior” stimulus) upon the BOLD fMRI response to the “subsequent” stimulus, averaged across the three participants. The carry-over effect of the chromatic L–M and S cone directed stimuli were combined into a single chromatic measure. The white asterisks indicate the cells in which the average modulatory effect differed from zero by more than two times the standard error of the mean of that measure across participants. Extended Figure 6-1 presents the separate carry-over matrices for each participant.

For achromatic stimuli, trials that presented no or low-frequency flicker (0, 2, 4 Hz) resulted in a larger than average BOLD fMRI response in the subsequent trial. Conversely, presentation of more rapid (8, 64 Hz) flicker tended to produce attenuation of response in the subsequent trial. A more complex pattern of effects was seen for 16 and 32 Hz flicker, which enhanced the responses of some, but not all, subsequent stimuli. A similar, albeit smaller, pattern of carry-over effects was seen for chromatic modulations. These general features can be observed in the separate carry-over matrices for each participant (Figure 6-1).

### Distributed patterns of response evoked by achromatic and chromatic flicker

We find that 64 Hz achromatic flicker produces only a minimal response within area V1 (Figure 4). Despite this, our measure of carry-over effects showed that 64 Hz achromatic flicker reliably suppressed the response to subsequent stimuli. We considered that our stimuli might induce other subtle changes in neural activity not revealed in the mean amplitude of response across the cortex. In split-half partitions of the data across acquisitions we examined the similarity of multi-vertex patterns of BOLD fMRI response evoked by the stimuli (relative to the 0 Hz stimulus) within area V1. Figure 7 presents the pattern similarity matrices we derived across stimulus frequency for the achromatic and chromatic stimuli, expressed as Pearson correlation between the split-halves of the data, and averaged across participants. The matrices derived from the separate subjects are in good agreement (Figure 7-1). The presence of high correlation values along the diagonal indicates that there are reproducible patterns of response across V1 evoked by the stimuli. Positive correlations in the off-diagonal cells indicate that there is some similarity in the patterns evoked by other flicker frequencies, rendering them confusable. Note that, while the mean amplitude of BOLD fMRI response is similar for 8 and 32 Hz LMS flicker (Figure 3a), the evoked patterns are negatively correlated. The pattern of response evoked by 64 Hz achromatic flicker is both highly reproducible and distinct, while this is not the case for 64 Hz chromatic flicker. The presence of 64 Hz achromatic flicker could be decoded with effectively 100% accuracy at every eccentricity within area V1 (Figure 7-2).

**Figure 7.**
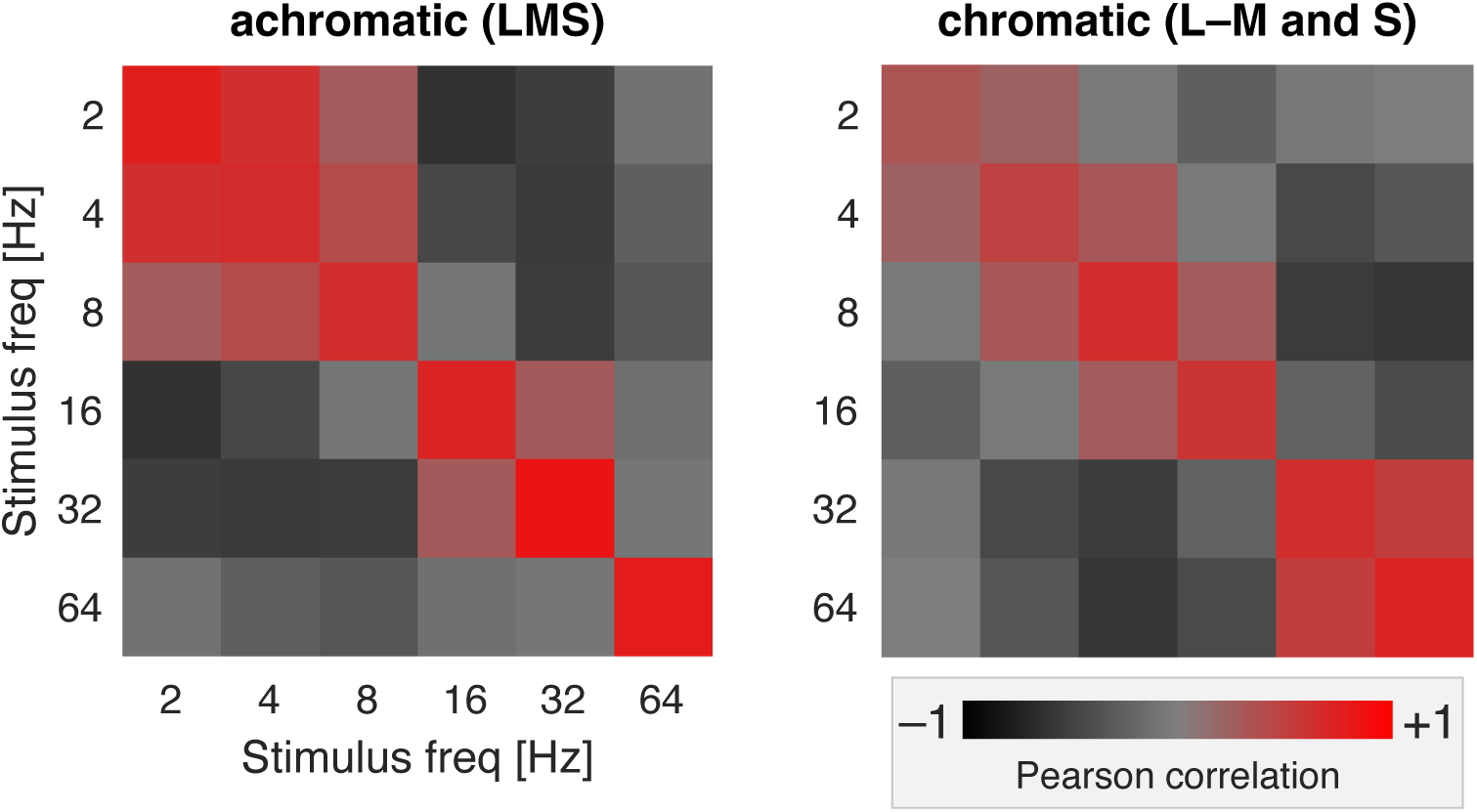
Multi-vertex pattern similarity in V1. Each matrix presents the split-half correlation of the multi-vertex pattern in area V1 evoked by stimulation with one frequency, as compared to stimulation with other frequencies. Separate matrices were derived for stimulation with achromatic (right) and chromatic (left) flicker, averaged across the three participants. The matrices are diagonally symmetric. Cells with high off-diagonal correlation indicate ‘areas of confusion’ where stimuli of different frequencies are not clearly distinguishable. This was most notable for low achromatic frequencies and high chromatic frequencies. Extended Figure 7-1 contains the separate pattern similarity matrices for each participant. Extended Figure 7-2 presents decoding performance for 64 Hz flicker as a function of eccentricity.

### Effect of the attention task

Participants performed an attention task during scanning in which they monitored for and responded to an infrequent event in which the stimulus field dimmed for 250ms. We modeled and removed the effect of this attention task in the fMRI data. As a check on the outcome of this procedure, we created a map of the effect of the attention event upon the BOLD fMRI signal within posterior cortex, averaged across participants (Figure 8). The most reliable responses were found throughout area V1. Additional areas of response were found more anteriorly, in the superior temporal and parietal lobes, but not in the early extra-striate visual areas (compare with the visual areas illustrated in Figure 5a). We further confirmed that the shape of temporal sensitivity function within area V1 was the same when constructed using trials that did, or did not, include an attention event (Figure 8-1). These findings suggest that our design was effective in capturing the variance introduced by the attention task.

**Figure 8.**
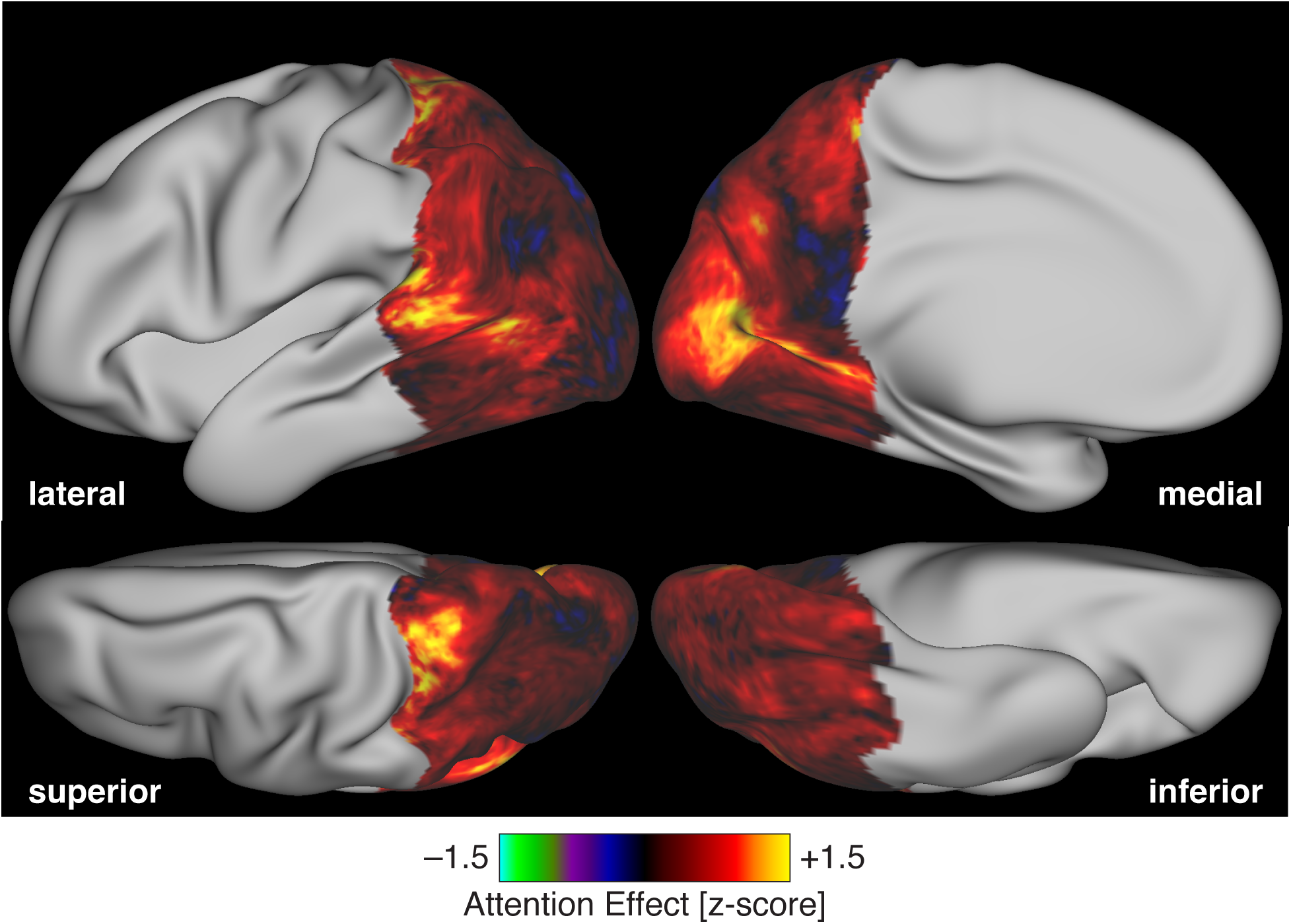
Map of the effect of attention events. The effect of the attention events, expressed as the mean BOLD fMRI signal evoked by attention events, divided by the standard deviation of the evoked response across acquisitions. The resulting z-maps from each participant were then averaged across participants. The entire extent of area V1 exhibited a reliable response to these events. Temporal sensitivity functions derived from the stimulus trials that did or did not include an attention event were quite similar for the three participants, and are shown in Figure 8-1.

### Comparison of V1 responses to predicted retinal signals

To better understand the transformation of signals from the retina to the cortex, we generated a prediction of integrated retinal output to our stimuli, based upon prior measurements of macaque RGC responses (Solomon et al., 2002; Solomon, 2005). In Figure 9, the filled circles and lines correspond to the predicted retinal response, while less-saturated squares, triangles, and circles are the data measured from our three participants within area V1 across eccentricity.

**Figure 9.**
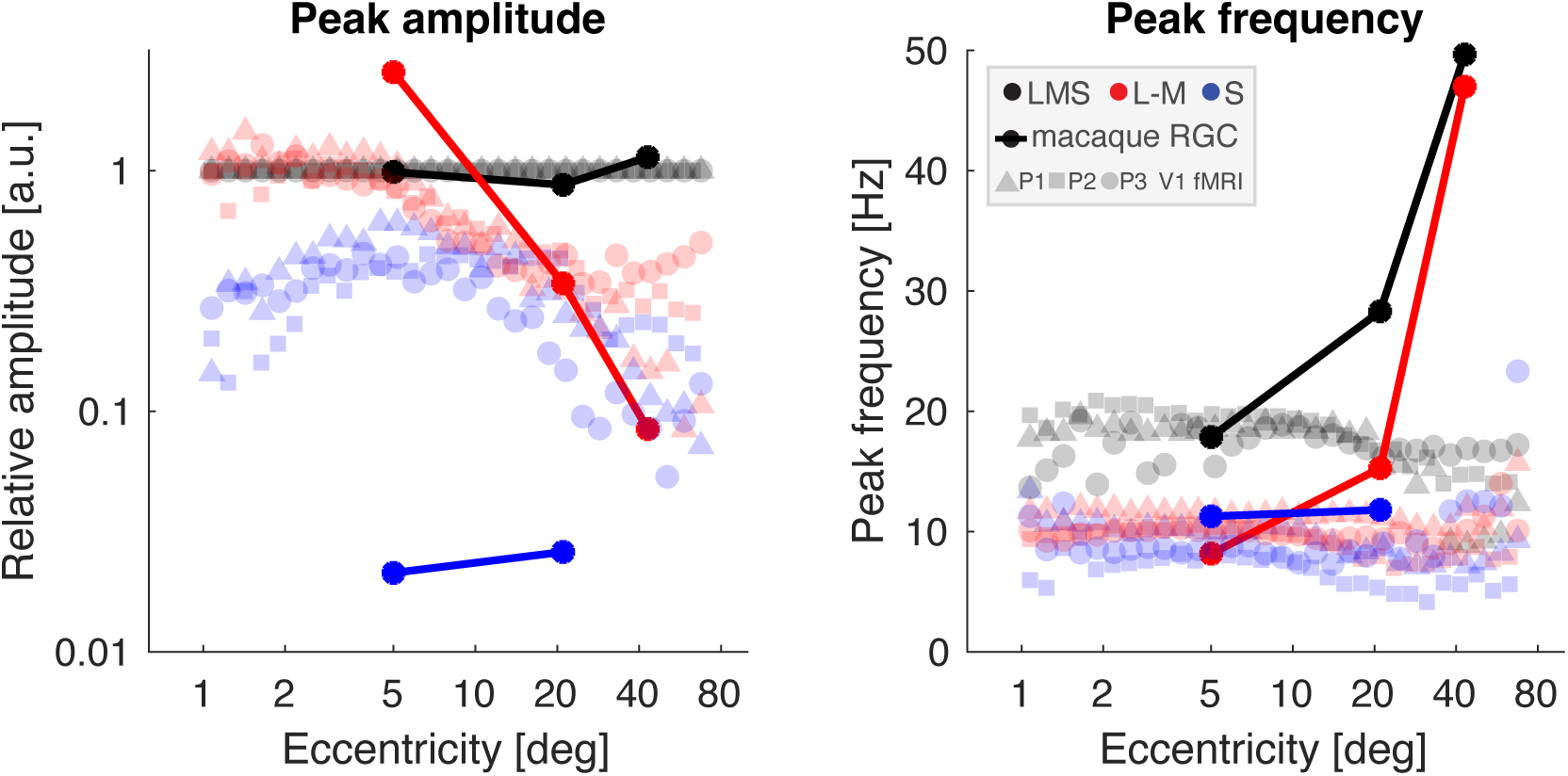
Predicted retinal signals compared to V1 response. Peak amplitude (left) and peak temporal frequency (right) of the V1 fMRI data from all three participants (triangles, squares, and circles) compared to predicted retinal output (filled circles and lines) based upon physiologic recordings of macaque RGCs, integrated across the total cell population present at a given eccentricity. Peak amplitude is expressed relative to responses to LMS-directed stimuli. For the retina model, normalization is to the average response evoked by LMS stimuli across eccentricity; for the V1 data, normalization is performed at each of 30 eccentricity bins.

The predicted amplitude of retinal response to LMS stimulation varies only a small amount among the three eccentricity locations for which measurements are available. This is in contrast to the retinal response to L–M stimulation, which is initially greater than the response to LMS, but drops by an order of magnitude into the visual periphery, as the combined consequence of both a relative reduction in the midget RGC class and the reduced response per cell, perhaps due to the effects of random cone wiring (Wool et al., 2018). Finally, the predicted response to S-directed stimulation is markedly smaller due to the sparse population of small bistratified cells that are presumed to carry this signal.

We may compare the model output to the fMRI data, normalized at each eccentricity location by the response to LMS stimulation. Similar to the retinal prediction, the relative response to L–M stimulation drops steadily beyond 5-10° eccentricity, although the V1 response is smaller than might be predicted in central vision. The amplitude of response to S-directed stimuli within V1 is markedly larger than predicted by the retinal model, rising further towards 5-10° eccentricity, then falling in concert with the response to L–M.

The retinal model predicts the temporal frequency that would elicit the peak response for each post-receptoral direction. At 5° eccentricity, there was rough agreement between the retinal model and the fMRI data. At greater eccentricities, however, retinal responses to LMS and L–M stimuli were predicted to shift to ever higher peak temporal frequencies, in marked contrast to the properties of the fMRI data.

## Discussion

Our study provides detailed temporal sensitivity measurements across subcortical and cortical regions in response to supra-threshold stimulation of the three canonical, post-receptoral visual channels. These results also characterize a fundamental aspect of human cortical visual function across eccentricity. Many of our findings replicate known properties of the visual system, including a relative amplification of chromatic signals between the LGN and area V1 (Cottaris and De Valois, 1998; Mullen et al., 2008), filtering of high temporal frequencies between these neural stages (Horwitz, 2021a), a systematic difference between chromatic and achromatic channels in peak temporal sensitivity (Swanson et al., 1987; Spitschan et al., 2016), and faster peak temporal sensitivity in dorsal extra-striate cortex (Liu and Wandell, 2005; Spitschan et al., 2016). Contrary to what might be expected from the properties of the retina and threshold psychophysical measurements, however, we do not observe an increase in peak temporal sensitivity to achromatic stimuli in the visual periphery.

### Comparison to prior neuro-imaging studies

There have been prior studies of the temporal sensitivity for achromatic and chromatic modulations within primary visual cortex (Liu and Wandell, 2005; D’Souza et al., 2011) and extra-striate cortex (Liu and Wandell, 2005; Spitschan et al., 2016), but these experiments did not examine responses across eccentricity. Chai and colleagues (2019) measured BOLD fMRI responses to a spatially structured, 15° radial eccentricity, achromatic checkerboard pattern that flickered between 1 and 40 Hz. Their parcellation of cortex by temporal sensitivity is similar in appearance to our temporal sensitivity map (Figure 5a). Kim and colleagues (Kim et al., 2023) developed a spatio-temporal receptive field model and analyzed fMRI data collected in response to drifting bars that contained multi-colored images. They find that a temporal integration parameter increases across the first 12 degrees of V1 eccentricity, although there is not a straightforward relationship between their model and our measurements.

Himmelberg and Wade (2019) reported that V1 response contrast sensitivity for a 20 Hz achromatic grating was greater at 6-10° eccentricity as compared to the fovea. As we did not measure a neural contrast response function for our stimuli, we are not able to directly compare this prior result to our own. Additionally, temporal sensitivity varies substantially with luminance and spatial frequency (Watson, 1986), and these stimulus properties differ between the current study and that of Himmelberg and Wade. We measured responses to a spatially uniform field at high retinal illuminance and leveraged high-dimensional spectral control to equate stimulus contrast across eccentricity. An advantage of this approach is that we need not contend with the effect of variation of acuity across the visual field (Virsu et al., 1982).

### Post-retinal modification of temporal sensitivity

Comparison of our measurements from area V1 to a model of retinal response suggests that there is extensive post-retinal modification of visual signals across eccentricity. Variation in response amplitude in V1 across stimulus directions is a log unit less than predicted to arise in the retina. There is also a notable difference between predicted and observed temporal sensitivity with eccentricity: while the visual cortex data at 5° eccentricity is consistent with the retinal prediction, at further eccentricities cortical responses peak at far lower stimulation frequencies than predicted. It appears that post-retinal modification between the retina and V1 normalizes responses across eccentricity and post-receptoral channel, modulating the relative amplitude of chromatic responses and slowing peak temporal sensitivity towards the periphery.

It is likely that this normalization arises with successive filtering at the LGN and V1 stages. Due to stronger surround inputs, LGN neurons exhibit a more transient response to step stimuli and a larger amplitude “rebound”, converting low-pass RGC responses to a more band-pass form (Ghodrati et al., 2017). Additionally, at high stimulus contrast levels, “bursty” LGN neurons exhibit low-pass filtering, with a transfer ratio that declines steadily above approximately 5 Hz (Mukherjee and Kaplan, 1995). Further low-pass filtering takes place at the geniculo-cortical synapse (Horwitz, 2021b). It seems that post-retinal temporal filtering would need to vary by eccentricity to account for the normalized V1 signal we observe. Our data do not speak to the degree to which this eccentricity-dependent normalization takes place within the LGN, V1 or at both levels.

We might consider alternative explanations for the relative stability of temporal frequency sensitivity that we observe across eccentricity in area V1. We derived temporal sensitivity measures from simultaneously stimulated eccentricities. It is possible that lateral or hierarchical influences between neurons across eccentricity locations cause homogenization of responses (Nassi et al., 2014). Alternatively, our stimulus might have been ineffective at driving neurons that have both narrower spatial tuning, and which vary in temporal tuning across eccentricity (Reinhard and Münch, 2021). In either case, measurements made with spatially restricted stimuli that produce isolated stimulation at different eccentricities might produce different findings.

We designed our stimuli to be spatially uniform, including accounting for variation in spectral sensitivity of the cone fundamentals with eccentricity (Figure 1-1). Nonetheless, we may be concerned that stimulus contrast or background luminance declined into the periphery, causing a decrease in temporal sensitivity (Snowden et al., 1995). The absolute amplitude of BOLD fMRI response measured for all three stimulus directions decreased beyond approximately 10° eccentricity, potentially supporting this concern. We are reassured, however, that this effect is unlikely to account for the slowing of temporal sensitivity for the luminance channel, as the amplitude of response in the far periphery is equal to or greater than that found at the fovea, despite a slowing of the TSF. In future studies, measurement of temporal sensitivity at varying contrast levels could be used to better address this concern. This would address a potential discrepancy between our findings and a prior measure of temporal sensitivity of ventral visual cortex (Liu and Wandell, 2005), and potentially disambiguate the effects of non-neural influences upon the BOLD fMRI response (Kurzawski et al., 2021).

### Relation to psychophysical measures at and above perceptual threshold

The cortical temporal sensitivity functions we have measured are in good agreement with known properties of human perception. This includes both the general form of the responses—which are well-fit by the Watson difference-of-exponentials model—and the variation in peak temporal sensitivity between the chromatic and achromatic channels.

Perceptual temporal sensitivity across visual field eccentricity has been well studied for the detection of high-frequency achromatic flicker. Specifically, the critical flicker fusion (CFF) frequency is a measure of highest frequency stimulus that can be distinguished from a static background. There is evidence that people are able to perceive higher frequency achromatic flicker in the periphery of their visual field (Hartmann et al., 1979; Kelly, 1984; Tyler and Hamer, 1993). Recent work suggests that this improvement in threshold detection is due to variation in the signal:noise ratio with eccentricity, as opposed to a change in the temporal sensitivity function itself (Rider et al., 2021).

Two of us (P1 and P3) exhibited a significant response to 64 Hz achromatic flicker in the V1 representation of the mid-peripheral visual field (8 – 16°), but not at foveal locations (Figures 4, 4-1). These two participants therefore demonstrated improved temporal sensitivity in the visual periphery in the sense of the highest frequency for which a non-zero response could be detected. Anecdotally, these observers perceived a faint, shimmering flicker in an annular zone of their peripheral vision during 64 Hz LMS flicker. We also measured the distributed neural patterns evoked by our stimuli. For all three observers there was a stable and distinct pattern of V1 response evoked by 64 Hz LMS stimulation, and this pattern was less reliable for chromatic flicker (Figure 7-1). These findings provide evidence of subtle V1 cortical responses to high frequency achromatic flicker, as has been reported previously (Williams et al., 2004).

Similar to our neuroimaging study, some perceptual studies of supra-threshold flicker perception have found equivalent performance at the fovea and periphery. Temporal frequency discrimination using high-contrast achromatic flicker is quite similar at the fovea and at 30° eccentricity (Waugh and Hess, 1994). Also, in a contrast-matching experiment, high-contrast stimuli that targeted the LMS, L–M, and S post-receptoral channels all appeared perceptually equivalent at the fovea and at 12° eccentricity (Jiang et al., 2022). The results of this study match our anecdotal experience, which is that the low-temporal frequency chromatic stimuli were perceived as fully saturated throughout the stimulus field.

### Neural adaptation

Prolonged presentation of a stimulus can modify the neural response to subsequent stimuli. These effects can be measured at levels throughout the visual hierarchy, including within the RGCs (Solomon et al., 2004). We measured the “carry-over” effect of stimulus history in our data. A reliable finding was that V1 neural responses were larger when preceded by stimuli that presented no or slow flicker, suggesting a recovery from habituation. Conversely, presentation of 64 Hz flicker caused a reduction in response to subsequent stimuli. This adaptive effect arose despite the absence of a substantial average neural response within area V1. This circumstance is consistent with a reduction in RGC firing, and with the finding of perceptual adaptation effects induced by flickering stimuli that are above the fusion frequency (Shady et al., 2004). Carry-over effects were more pronounced with achromatic as compared to chromatic stimuli, again consistent with the response of RGCs (Solomon et al., 2004).

There was a more complex pattern of carry-over effects following 16 and 32 Hz stimulation. We are unable to offer a simple account of these findings. Notably, we experienced robust, spatially structured entopic percepts while viewing these stimuli, consistent with prior reports (Purkyně, 1818). It is possible that the sequential experience of these phenomena interact in a complex manner.

Our use of a counter-balanced ordering of flicker frequency insulates our measurement of the “direct” effect of the stimulus from first-order carry-over effects (Aguirre, 2007). There are other forms of dynamic neural response, however, that may have influenced our measurements. For example, V1 neurons exhibit an exponential decay in response to prolonged visual stimulation (Vautin and Berkley, 1977). We measured the mean response to stimuli across a 12-second trial. Variation in the time constant of within-trial habituation of neural responses across this period could cause the temporal sensitivity functions we measure to differ from those that would be derived from the amplitude of the onset response (although trials were half-cosine windowed in an effort to reduce such an effect). Improvements in trial design, temporal resolution of the measure, and/or modeling approach (Groen et al., 2022) may be needed to address this aspect of the neural response.

### Differences and aberrations

Our primary findings are supported by highly similar results across our three observers; it is quite likely that these findings are representative of the general population (Laurens, 2022). There are some aspects of our results, however, that differ across participants. We collected extensive within-participant measurements, allowing us to reject in many cases that these differences are due to within-participant measurement error. Consequently, these findings could reflect true individual differences, attributable either to trait or state. In this category are differences between the participants in the shape of V1 temporal sensitivity functions. P3 had a peak temporal sensitivity to LMS flicker that was consistently 5 Hz lower than observed in the other participants, and a bimodal distribution of peak sensitivity to LMS flicker at low eccentricities. The participants also differed in the width of their LMS sensitivity functions, the degree to which these functions rolled off at low frequencies, and (as discussed above) the presence of responses to 64 Hz LMS flicker. Future studies might attempt relate this variation in temporal sensitivity to perceptual performance and individual biological differences.

There are other aspects of our data that were reliable across participants, but for which we cannot offer a biological account. All three participants had points in the far periphery of V1 cortex that reliably responded to 64 Hz S-directed flicker, but to no other S flicker frequency. These vertices contributed to the non-zero values at 64 Hz in S-directed TSFs (Figure 4, 4-1; most prominent for P2) and the scattering of points of high temporal sensitivity in the visual field maps of V1 response (Figure 4-2). Similar points of cortical response were seen for the L–M directed stimuli, and these far-periphery responses contribute to an upturn of the temporal sensitivity values at the farthest eccentricities for the chromatic stimuli (Figure 4a, bottom row, rising median temporal sensitivity values beyond 40° eccentricity). It is possible that transverse chromatic aberration in the optics of the stimulus eye piece or the eye itself generated LMS contrast from the nominally iso-luminant chromatic stimuli in the far periphery, and that we are effectively measuring the response to 64 Hz luminance flicker at these points.

## Conclusions

This study provides the eccentricity dependence of the temporal sensitivity functions for the three cardinal post-receptoral channels within V1, and the spatially averaged sensitivity functions across multiple levels of the visual hierarchy. After the LGN, peak temporal sensitivity increases across the visual cortical hierarchy. We find that responses in primary visual cortex across post-receptoral channel and temporal frequency are substantially normalized relative to predicted signals from the retina.

## Author Contributions

Conceptualization: Manuel Spitschan, Andrew S. Bock, and Geoffrey K. Aguirre. Data curation: Huseyin O. Taskin. Formal analysis: Carlyn Patterson Gentile, Manuel Spitschan, and Geoffrey K. Aguirre. Funding acquisition: Geoffrey K. Aguirre. Investigation: Andrew S. Bock and Geoffrey K. Aguirre. Methodology: Geoffrey K. Aguirre. Project administration: Geoffrey K. Aguirre. Resources: Geoffrey K. Aguirre. Software: Carlyn Patterson Gentile, Manuel Spitschan, Huseyin O. Taskin, and Geoffrey K. Aguirre. Supervision: Geoffrey K. Aguirre. Validation: Manuel Spitschan and Geoffrey K. Aguirre. Visualization: Carlyn Patterson Gentile, Manuel Spitschan, and Geoffrey K. Aguirre. Writing - original draft: Carlyn Patterson Gentile and Geoffrey K. Aguirre. Writing - review & editing: Carlyn Patterson Gentile, Manuel Spitschan, Huseyin O. Taskin, Andrew S. Bock, and Geoffrey K. Aguirre.

## Acknowledgements

none.

## Abbreviations

LGN: Lateral geniculate nucleus
TSF: Temporal Sensitivity Function
RGC: Retinal ganglion cell
V1: Primary visual cortex

## EXTENDED DATA

**Figure 1-1.**
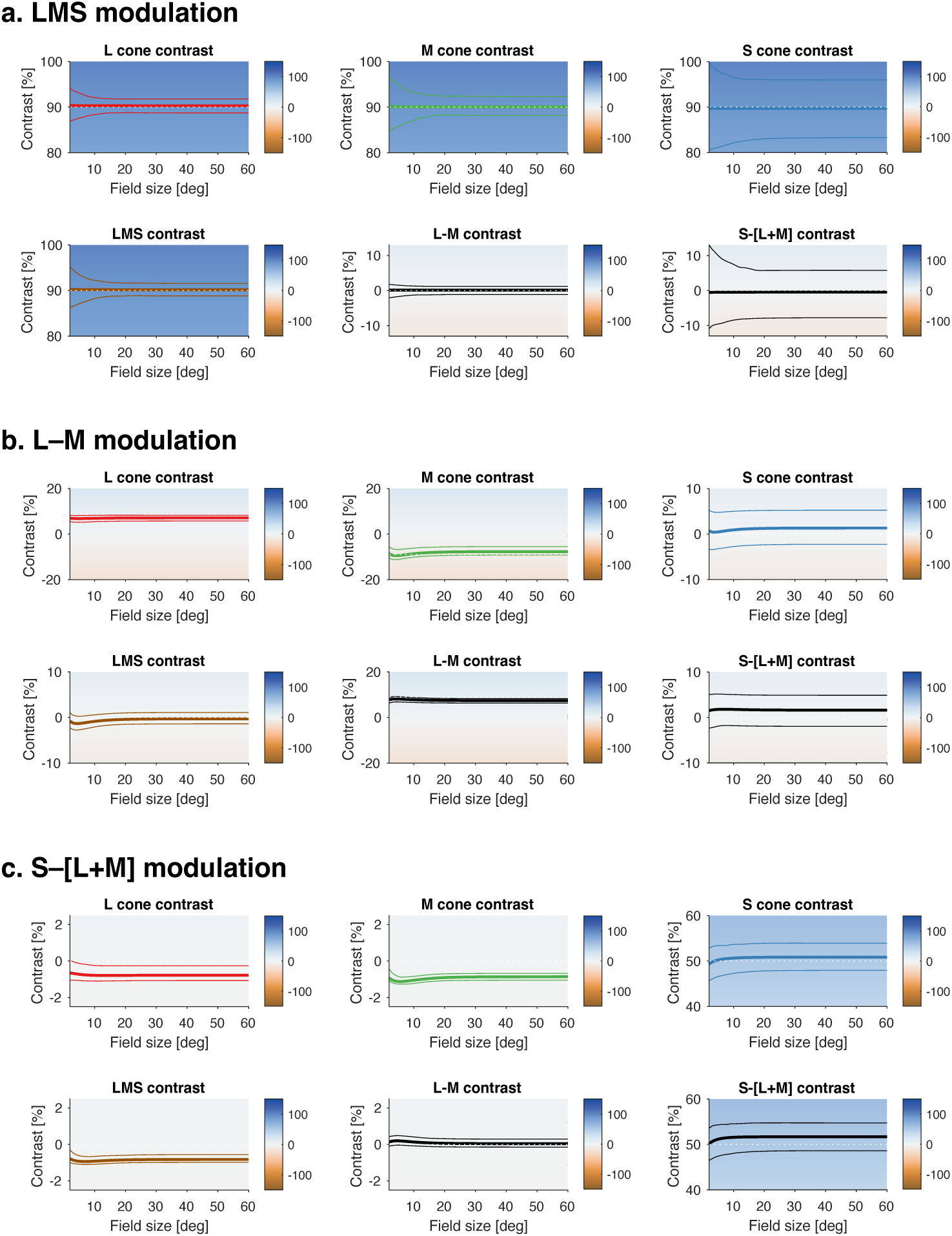
Validated and inadvertent stimulus contrast. Spectroradiometric validation measurements of the stimuli were made before the first scanning session. The contrast produced by these stimuli upon the cones and their post-receptoral combinations was calculated for a nominal, 32 year old observer as a function of eccentricity (the thick line in each plot). The effect of biological variation in the assumed properties of the photoreceptors and pre-receptoral filtering was estimated by Monte-Carlo simulation. The 95% confidence interval around the predicted contrast is given by the thin lines. Panels (a, b, c) correspond to the three stimulus modulations used in the experiment. The calculated contrast and confidence interval values at 2 and 30° is given in Extended Table 1-1.

**Figure 2-1.**
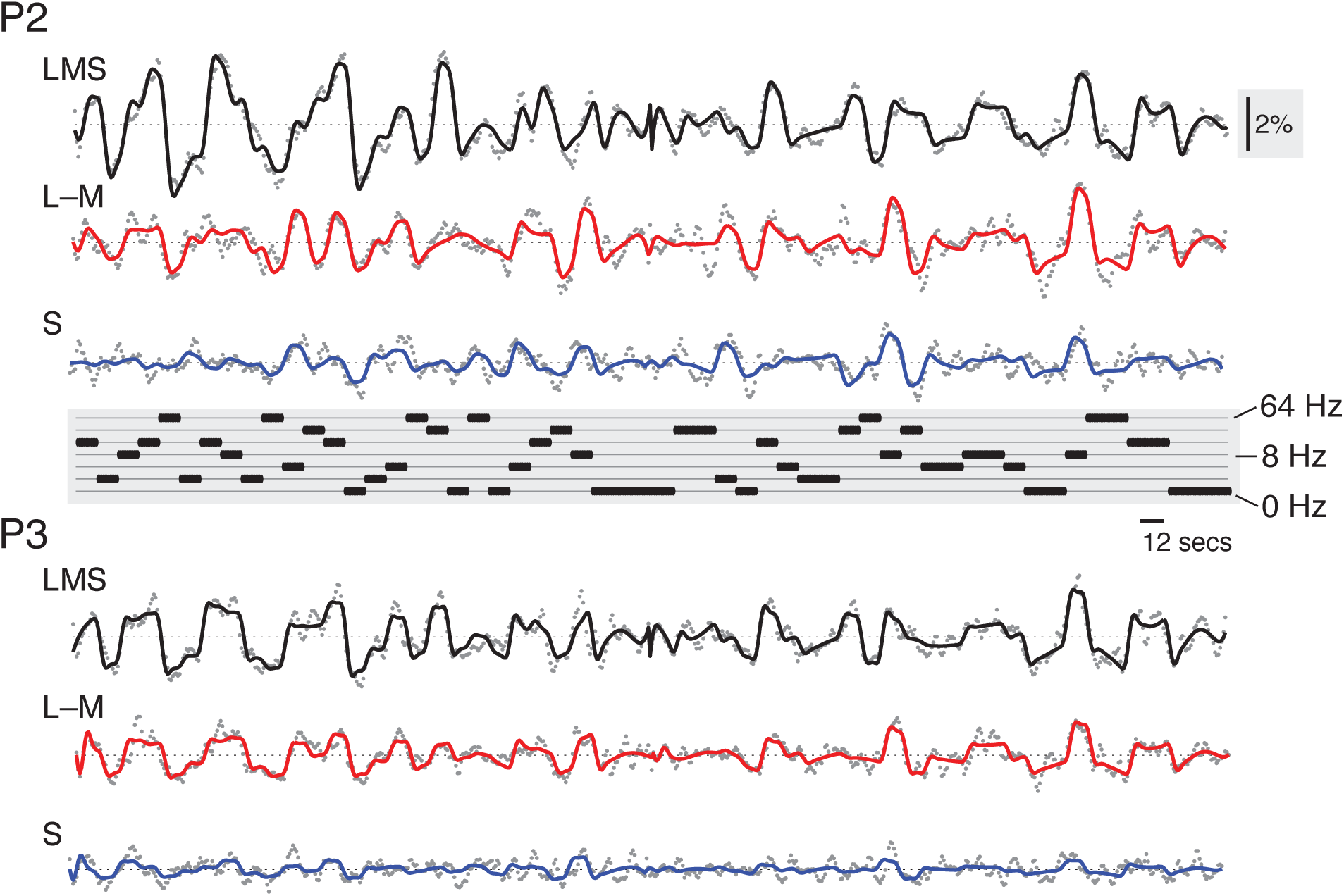
Fit to fMRI time series data for P2 and P3. The average BOLD fMRI time series (gray dots) from area V1, and across the acquisitions for each stimulus type (LMS, L–M, and S) for P2 and P3. The continuous line shows the fit of the time-series model. An inset, vertical scale bar on the right indicates a 2% BOLD fMRI signal response. An inset scale bar at center-right indicates a 12-second interval. The plot in the center provides the ordering of stimulus frequency.

**Figure 3-1.**
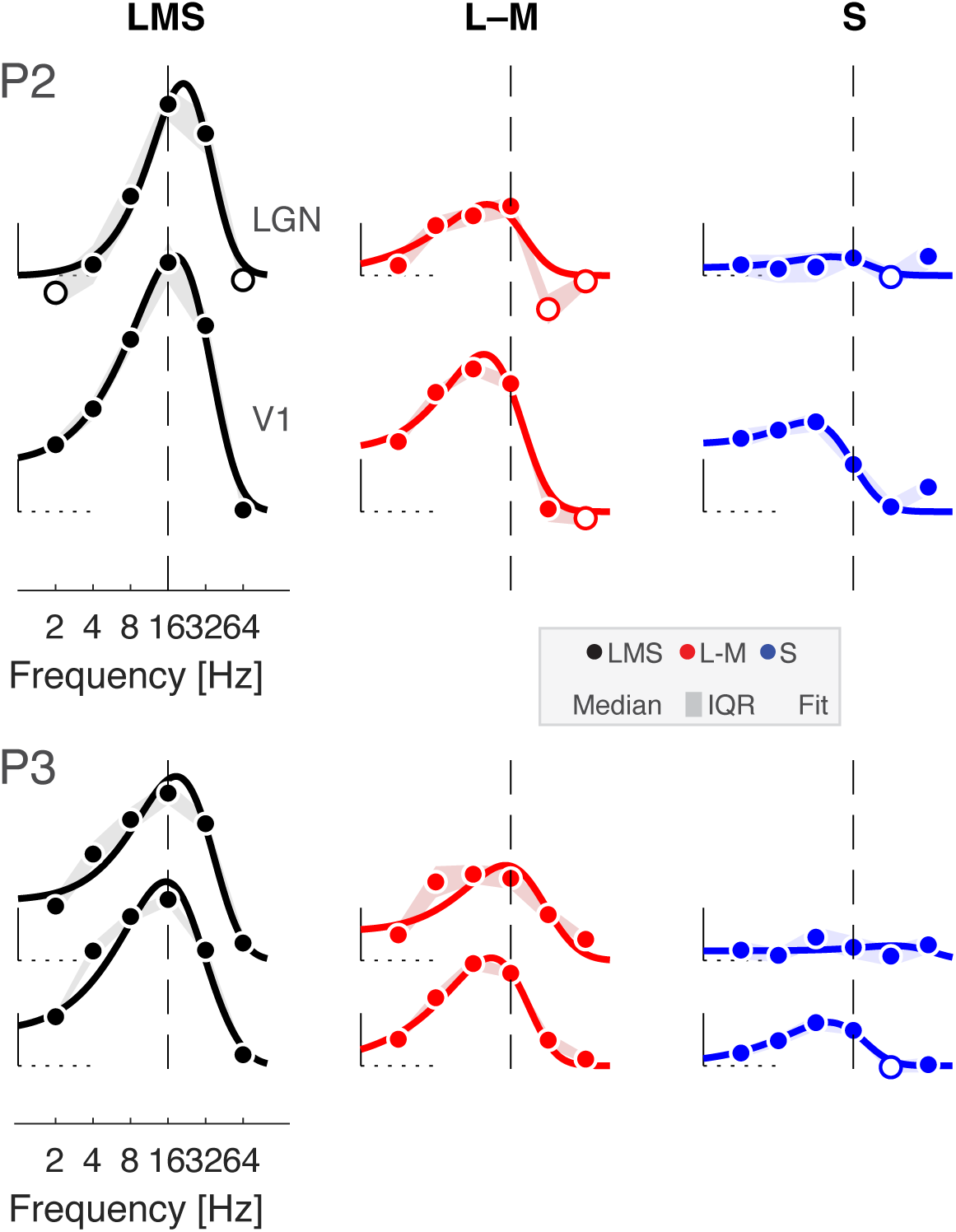
Temporal sensitivity to flicker in LGN and V1 for P2 and P3. LGN and V1 TSFs for stimuli that target the three post-receptoral channels are shown for P2 and P3, fit with the Watson difference-of-exponentials model. Data points represent the boot-strapped median across acquisitions, and shading provides the interquartile range. Open circles indicate points where the response was less than zero. A dashed reference line is positioned at 16 Hz in each plot, facilitating a comparison of the point of peak temporal sensitivity across plots.

**Figure 4-1.**
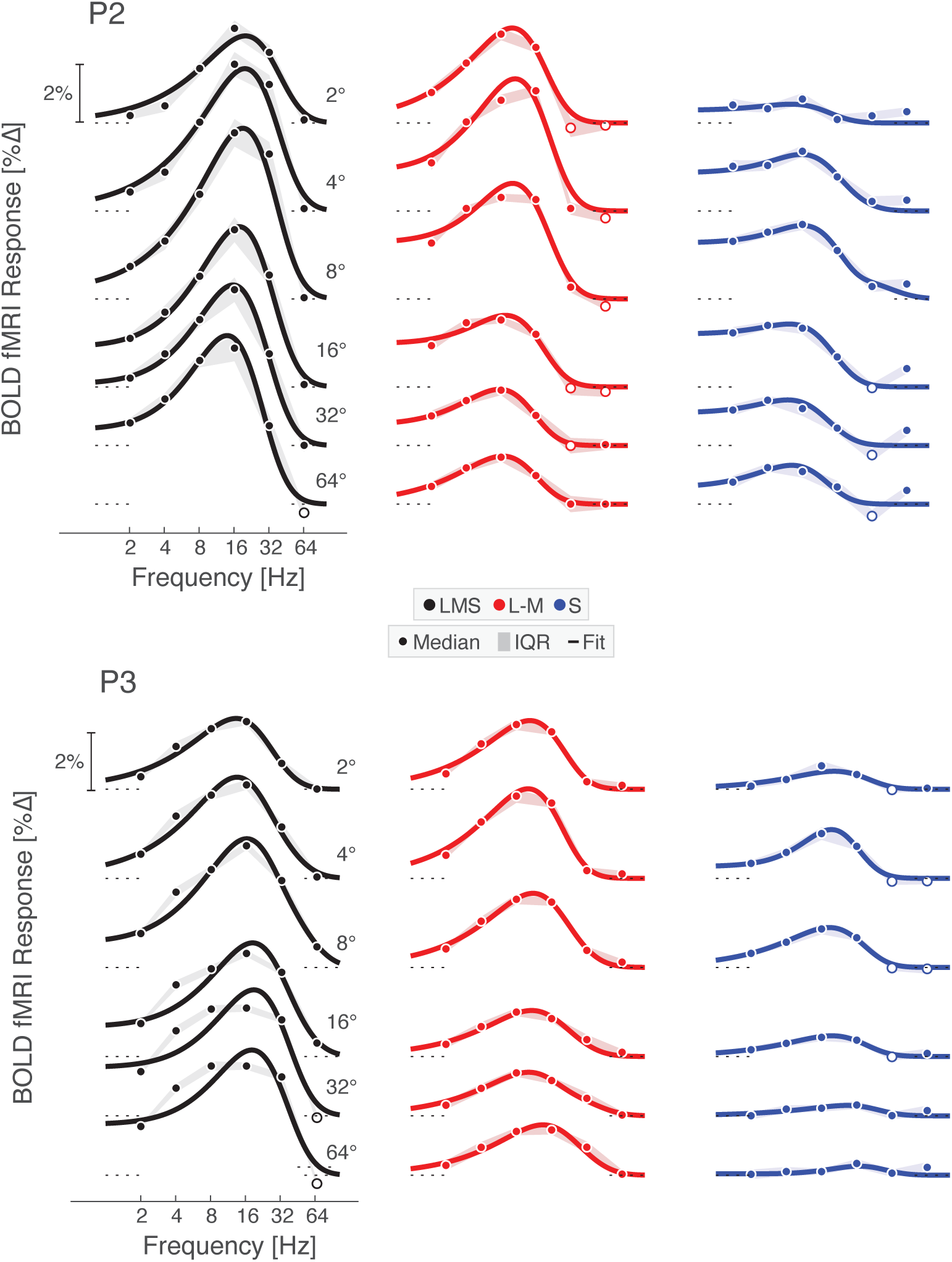
Temporal flicker sensitivity within area V1 across eccentricity. V1 TSFs at logarithmically spaced eccentricities across the three post-receptoral channels are shown for P2 and P3 with Watson temporal model fits. Data points represent the boot-strapped median across acquisitions, shading provides the interquartile range, and the line provides the Watson model fit. Open circles indicate points where the response was less than zero.

**Figure 4-2.**
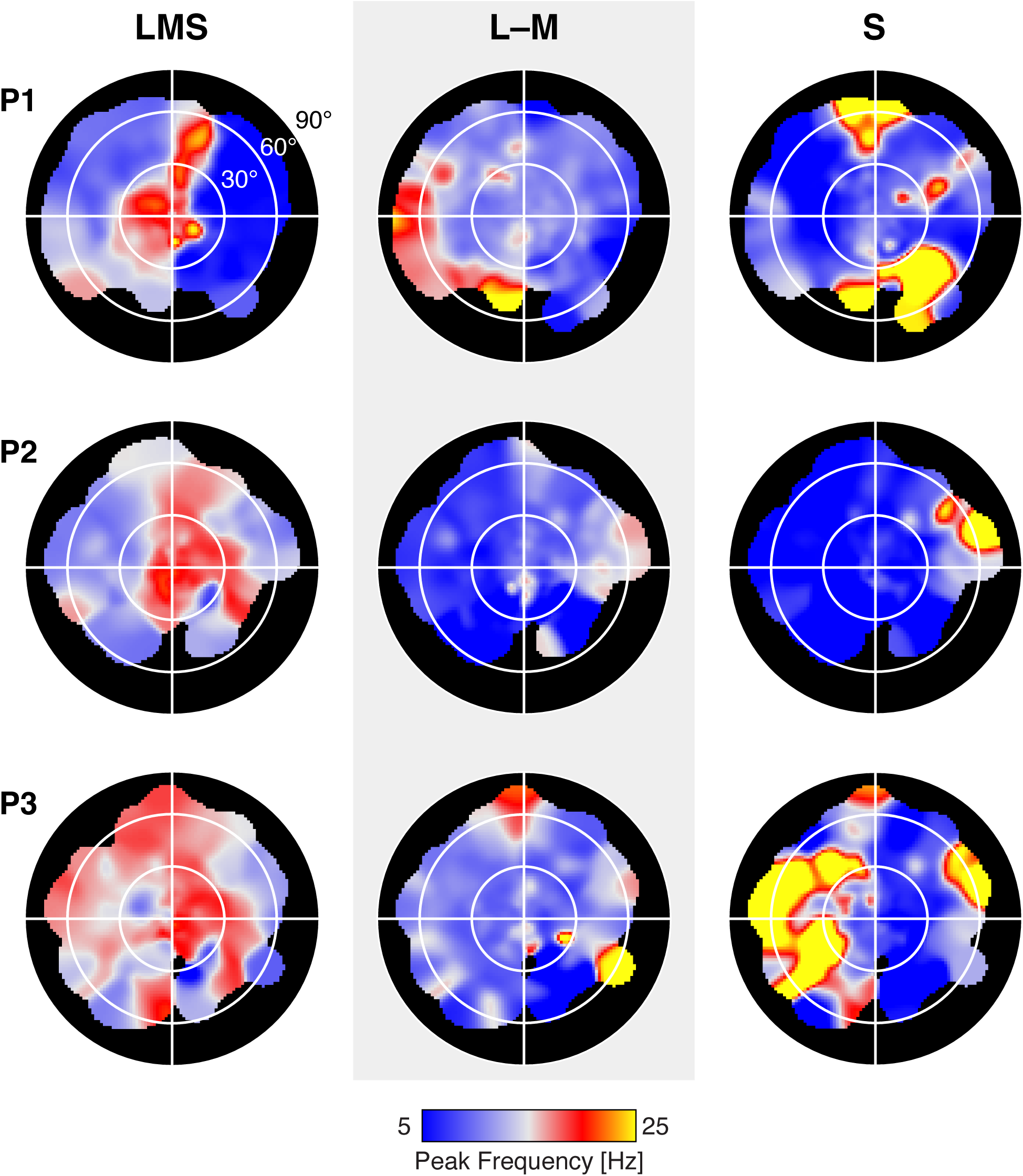
Visual field representation of peak temporal sensitivity. The peak temporal sensitivity for each stimulus direction was measured at each vertex within V1. The retinotopic mapping values of eccentricity, polar angle, and width of the population receptive field were then used to project cortical values to a visual field plot. Separate plots are shown for each post-receptoral stimulus direction and participant. Annular white lines, and the outer edge of the black plot area, indicate the 30, 60, and 90° eccentricity locations.

**Figure 6-1.**
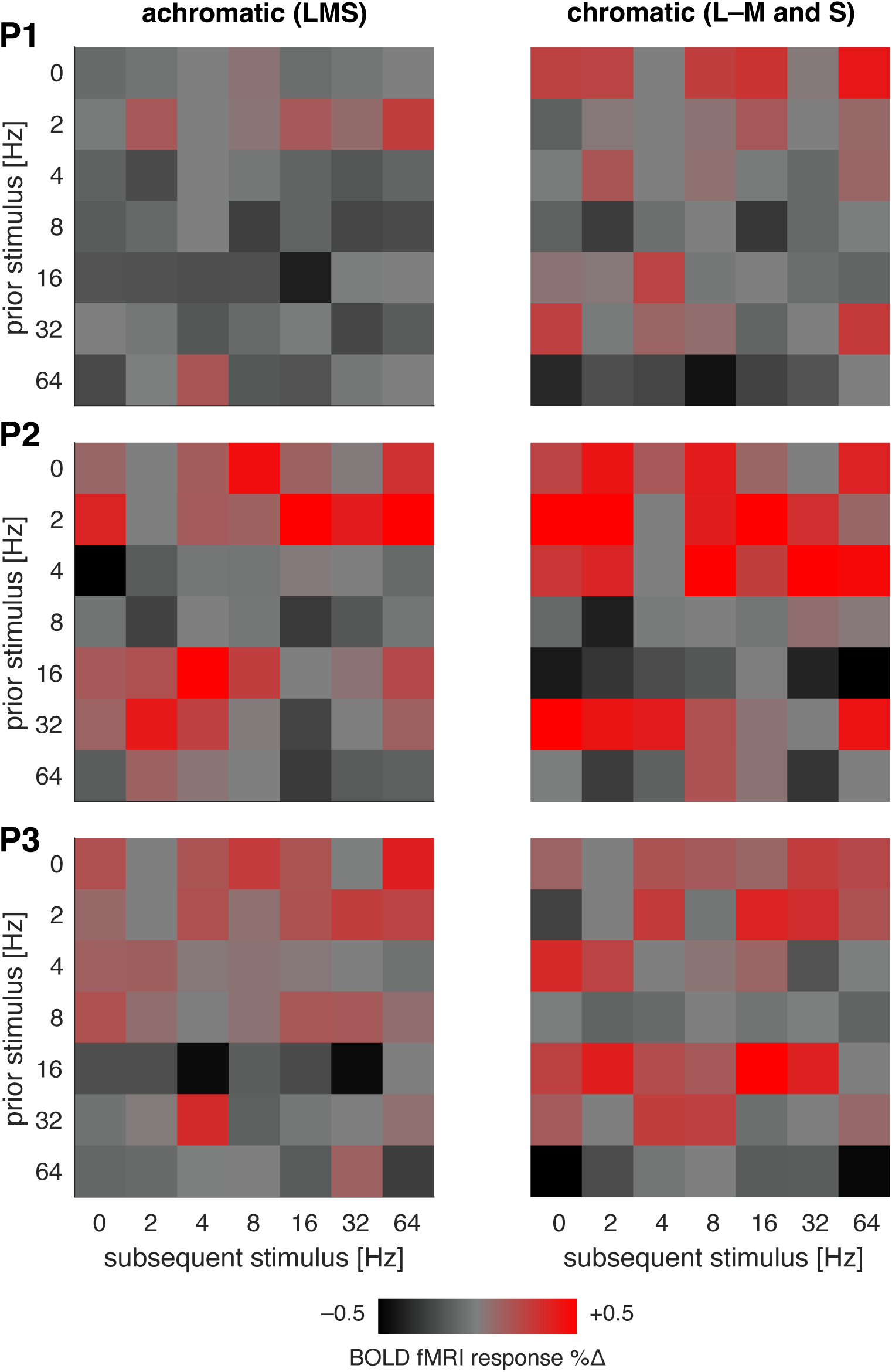
Carry-over adaptation effects in V1 by participant. Each matrix presents the modulatory effect of a given frequency of flicker stimulation (the “prior” stimulus) upon the BOLD fMRI response to the “subsequent” stimulus, with the data from each participant in rows. The carry-over effect of the L–M and S cone directed stimuli were combined into a single chromatic measure.

**Figure 7-1.**
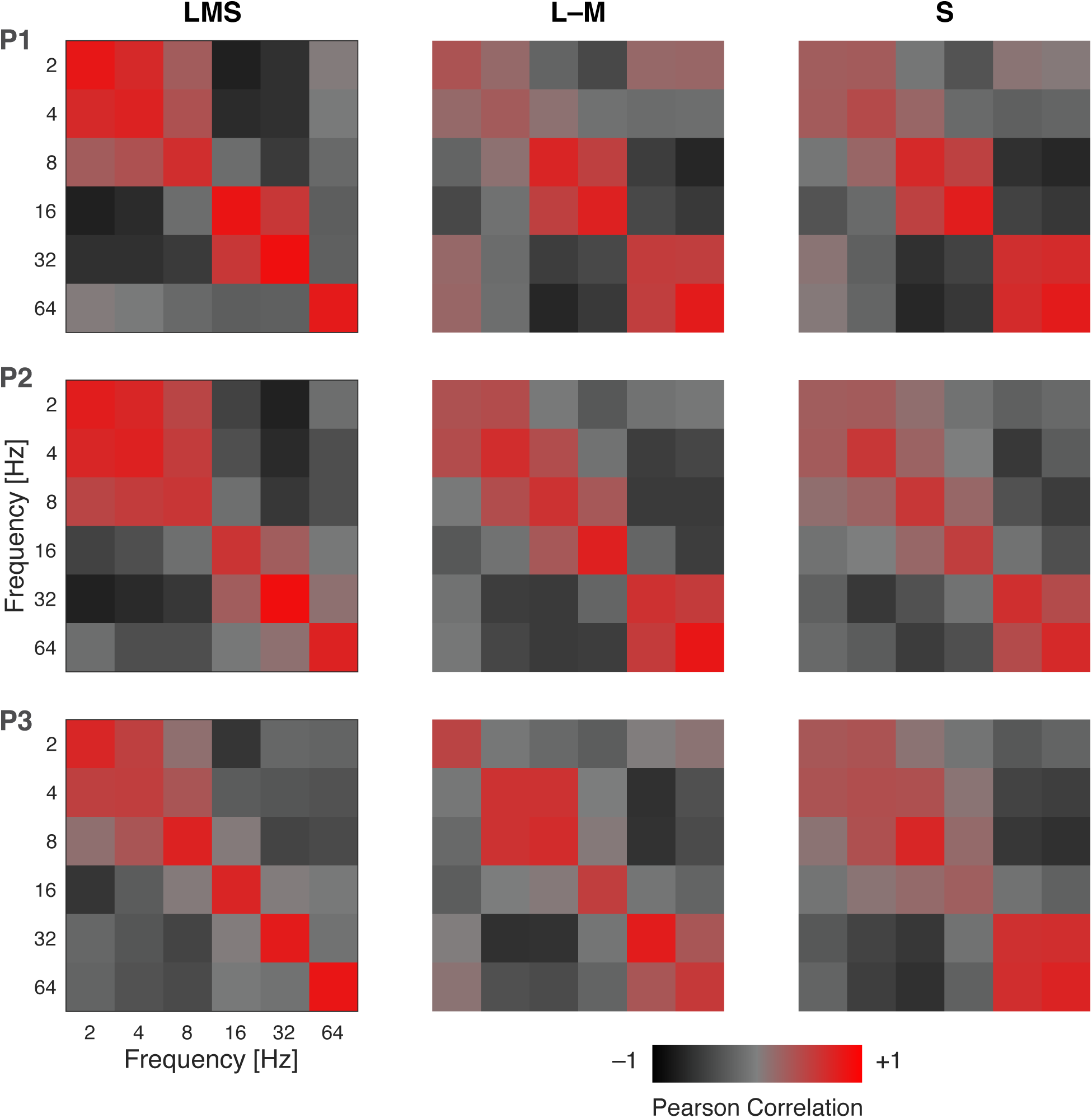
Multi-vertex pattern similarity in area V1 by participant. Each matrix presents the split-half correlation of the multi-vertex pattern in area V1 evoked by stimulation with one frequency, as compared to stimulation with other frequencies. Separate matrices were derived for stimulation in each of the post-receptoral directions; the two chromatic matrices were combined for presentation in Figure 7. The matrices are diagonally symmetric.

**Figure 7-2.**
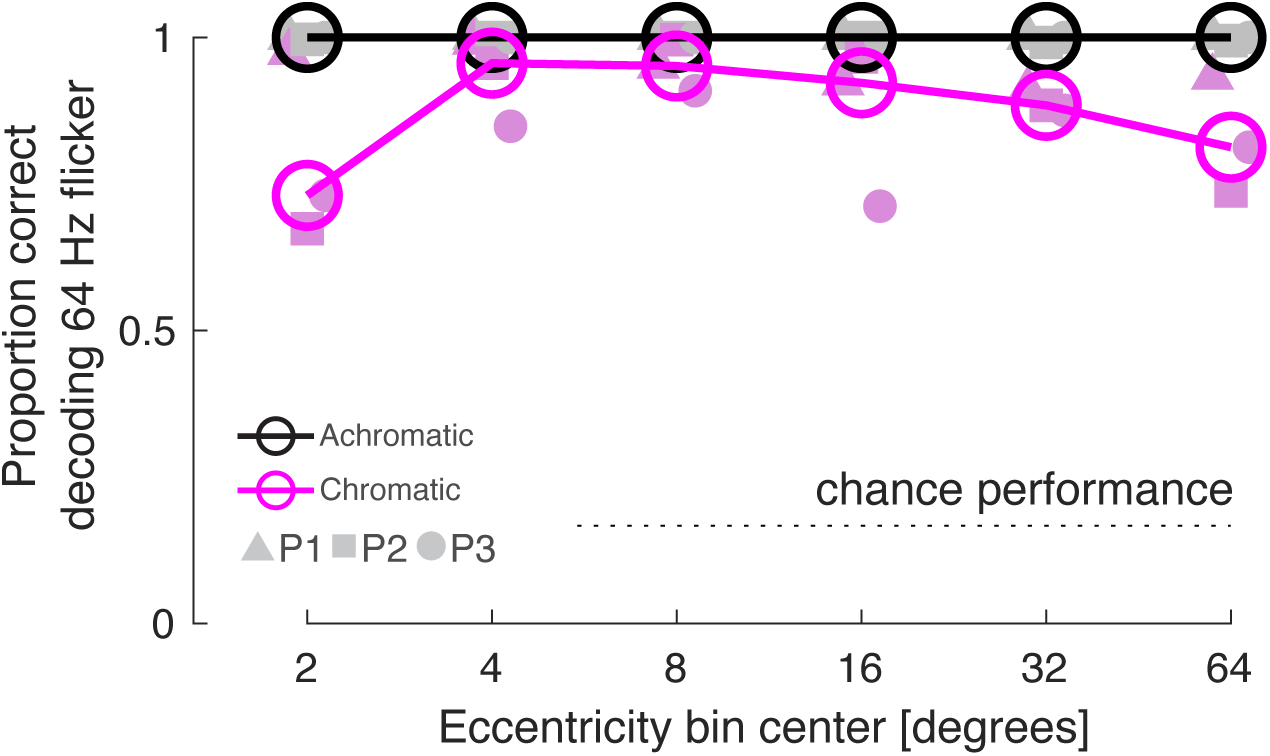
Decoding performance for 64 Hz flicker as a function of eccentricity. Across all split-half partitions of the imaging data we asked what multi-vertex pattern in the held-out data best matched the pattern generated by 64 Hz stimulation. The proportion of cases in which the pattern corresponding to 64 Hz flicker in the held-out data was selected as the best match provides the proportion of correct decoding (chance performance = 0.167). This calculation was performed for sets of vertices that were taken from bands of V1 cortex that varied in eccentricity. The decoding performance of individual participants for achromatic and chromatic 64 Hz flicker is indicated by the filled symbols, and the median performance across participants is given by the open circles and lines.

**Figure 8-1.**
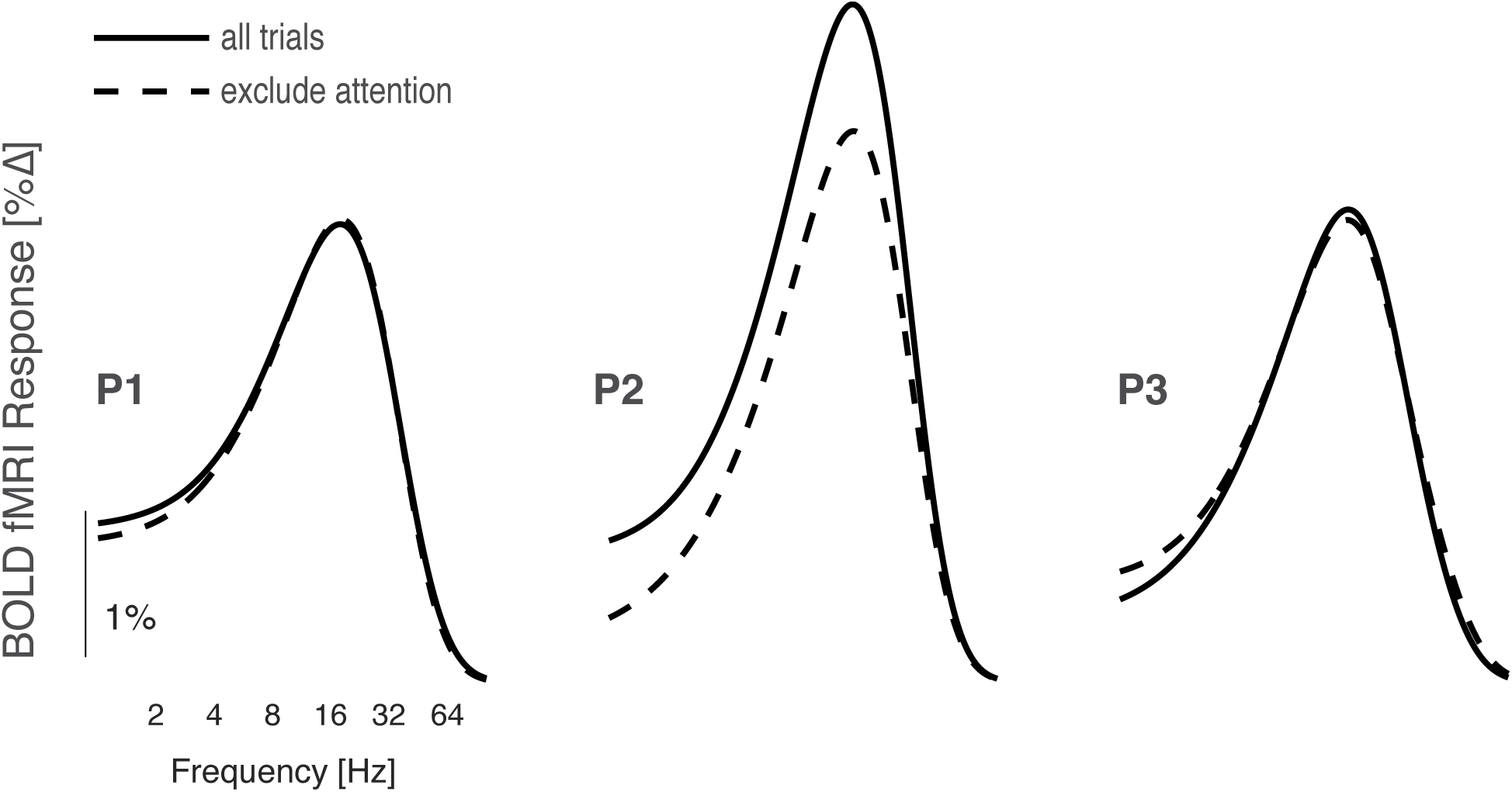
LMS temporal sensitivity functions that do or do not include attention trials. The average V1 temporal sensitivity for LMS flicker was derived using all of the available fMRI data (solid lines), and using just the ~2/3^rd^ of trials that did not include an attention event (dashed line). The resulting sensitivity functions are very similar in form, and for 2 of 3 participants matched in amplitude.

**Table 1-1.**
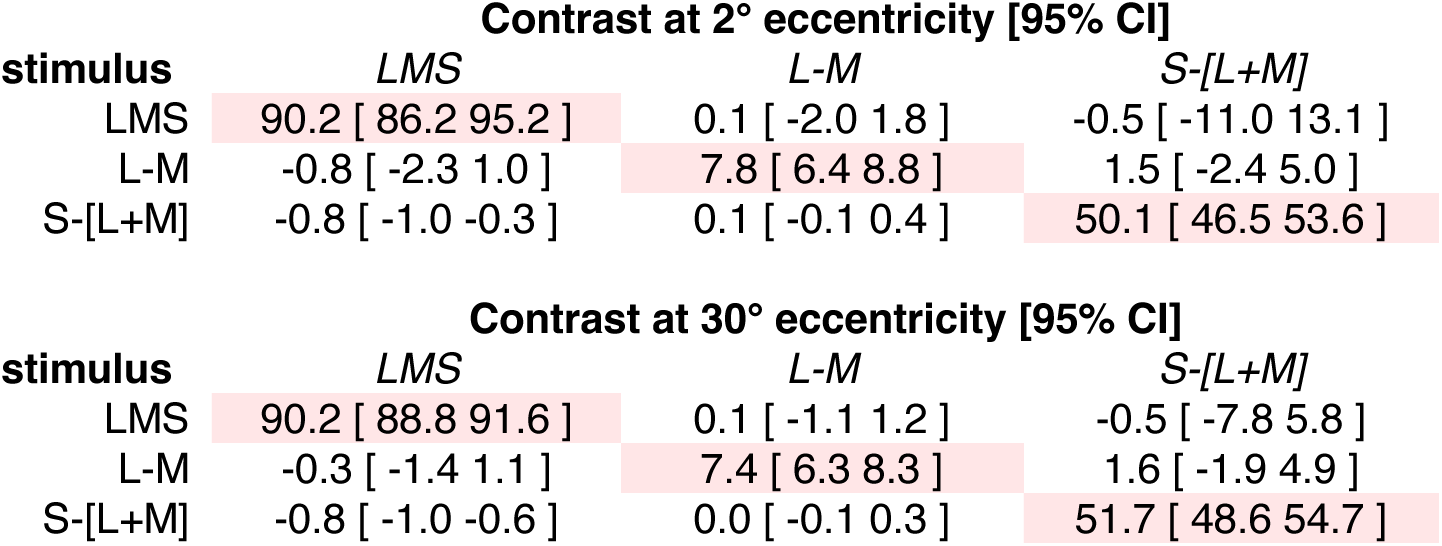
Stimulus contrast at 2 and 30 degrees eccentricity. The values presented here correspond to the thick and thin lines from Figure 1-1, at 2° and 30° on each plot that calculate contrast upon the post-receptoral channels. The cells that contain the contrast value for the targeted post-receptoral direction for each stimulus are indicated with red shading. Notably, there is minimal change in achieved contrast across a large range of eccentricity, and the off-diagonal contrast levels are uniformly low.

| Contrast at 2° eccentricity [95% CI] |  |  |  |
| --- | --- | --- | --- |
| stimulus | LMS | L-M | S-[L+M] |
| LMS | 90.2 [ 86.2 95.2 ] | 0.1 [ -2.0 1.8 ] | -0.5 [ -11.0 13.1 ] |
| L-M | -0.8 [ -2.3 1.0 ] | 7.8 [ 6.4 8.8 ] | 1.5 [ -2.4 5.0 ] |
| S-[L+M] | -0.8 [ -1.0 -0.3 ] | 0.1 [ -0.1 0.4 ] | 50.1 [ 46.5 53.6 ] |

**Table 1-1. Stimulus contrast at 2 and 30 degrees eccentricity.**
| Contrast at 30° eccentricity [95% CI] |  |  |  |
| --- | --- | --- | --- |
| stimulus | LMS | L-M | S-[L+M] |
| LMS | 90.2 [ 88.8 91.6 ] | 0.1 [ -1.1 1.2 ] | -0.5 [ -7.8 5.8 ] |
| L-M | -0.3 [ -1.4 1.1 ] | 7.4 [ 6.3 8.3 ] | 1.6 [ -1.9 4.9 ] |
| S-[L+M] | -0.8 [ -1.0 -0.6 ] | 0.0 [ -0.1 0.3 ] | 51.7 [ 48.6 54.7 ] |

